# Messenger RNA biomarker signatures for forensic body fluid identification revealed by targeted RNA sequencing

**DOI:** 10.1101/247312

**Authors:** E Hanson, S Ingold, C Haas, J Ballantyne

## Abstract

The recovery of a DNA profile from the perpetrator or victim in criminal investigations can provide valuable ‘source level’ information for investigators. However, a DNA profile does not reveal the circumstances by which biological material was transferred. Some contextual information can be obtained by a determination of the tissue or fluid source of origin of the biological material as it is potentially indicative of some behavioral activity on behalf of the individual that resulted in its transfer from the body. Here, we sought to improve upon established RNA based methods for body fluid identification by developing a targeted multiplexed next generation mRNA sequencing assay comprising a panel of approximately equal sized gene amplicons. The multiplexed biomarker panel includes several highly specific gene targets with the necessary specificity to definitively identify most forensically relevant biological fluids and tissues (blood, semen, saliva, vaginal secretions, menstrual blood and skin). In developing the biomarker panel we evaluated 66 gene targets, with a progressive iteration of testing target combinations that exhibited optimal sensitivity and specificity using a training set of forensically relevant body fluid samples. The current assay comprises 33 targets: 6 blood, 6 semen, 6 saliva, 4 vaginal secretions, 5 menstrual blood and 6 skin markers. We demonstrate the sensitivity and specificity of the assay and the ability to identify body fluids in single source and admixed stains. A 16 sample blind test was carried out by one lab with samples provided by the other participating lab. The blinded lab correctly identified the body fluids present in 15 of the samples with the major component identified in the 16^th^. Various classification methods are being investigated to permit inference of the body fluid/tissue in dried physiological stains. These include the percentage of reads in a sample that are due to each of the 6 tissues/body fluids tested and inter-sample differential gene expression revealed by agglomerative hierarchical clustering.

## 1. Introduction

Genetic identification of the donor of transferred biological traces deposited at the crime scene or on a person or implement using STR analysis is now routine practice worldwide [1]. This represents potentially crucial ‘source level’ information for investigators [2]. A DNA profile from the perpetrator does not however reveal the circumstances by which it got transferred. This contextual information (sometimes known as the ‘activity level’ in Cook and Evett’s classic 1998 paper [2]) is important for casework investigations because the deposition of the perpetrator’s biological material requires some behavioral activity on behalf of the individual that results in its transfer from the body. The consequences of different modes of transfer of the DNA profile may dramatically affect the investigation and prosecution of the crime. For example, a DNA profile from a victim that originates from skin versus the same DNA profile that originates from vaginal secretions may support social or sexual contact respectively. Thus tissue/body fluid sourcing of the DNA profile should be an important concern for, and service from, forensic genetics practitioners who are integral to the investigative team. The problem is that, up until the recent past, it was not possible to definitively identify many of the important body fluids of interest (e.g. vaginal secretions) and even now, it is not possible to conclusively link a DNA profile to a particular body fluid.

In order to overcome the limitations of currently used classical body fluid identification methods, the use of messenger RNA (mRNA) profiling has been proposed to supplant conventional methods for body fluid identification [3]. Terminally differentiated cells, whether they comprise blood monocytes or lymphocytes, ejaculated spermatozoa, epithelial cells lining the oral cavity or epidermal cells from the skin become such during a developmentally regulated program in which certain genes are turned off (i.e. transcriptionally silent) and turned on (i.e. are actively transcribed and translated into protein) [4]. Thus, a pattern of gene expression is produced that is unique to each cell type, which is evidenced by the presence as well as the relative abundance of specific mRNAs [4]. The type and abundance of specific targeted mRNAs, if determined, would then permit a definitive identification of the body fluid or tissue origin of forensic samples. This is the basis for mRNA profiling for body fluid identification. RNA profiling now offers the ability to identify most of the forensically relevant biological fluids using methods compatible with the current DNA analysis pipeline [3,5–10]. Despite the identification of numerous body fluid specific candidates there is some reluctance to utilize RNA profiling assays in the forensic community due to concerns over the perceived instability of RNA in biological samples, the unavailability of dedicated forensic commercial kits and the potential cost. Several studies have been conducted in order to assess the stability of RNA in dried forensic stains and have successfully demonstrated their suitability for use with aged and environmentally compromised forensic samples [11–16]. The recently published EDNAP collaborative exercises on mRNA profiling for body fluid identification further demonstrate a significant interest in mRNA profiling by the forensic community in Europe and around the world as well as the relative ease with which this technology could be implemented into forensic casework laboratories [17–22].

Up until present there have been three main methods developed for mRNA profiling: capillary electrophoresis (CE)-based analysis, quantitative RT-PCR and high resolution melt (HRM) analysis [5,23,24]. However an impediment to the implementation of mRNA profiling for body fluid identification has been the lack of a commercial product for any of these platforms but this may change with new technologies such as NGS. The advent and evolution of NGS in the past five years, which permits the massively parallel sequencing of millions of DNA fragments for a total read length per run ranging from 100 Mb-600 Gb, has revolutionized genomics [25]. Next generation high-throughput sequencing technology (NGS) is capable of the analysis of hundreds of loci useful for forensic genetic analysis [25] and has spawned the creation of a new operational paradigm, namely “forensic genomics”. Commercial forensic genomics kits for the massively parallel typing of autosomal STRs, Y-STRs, X-STRs, identity-SNPs, phenotypic-SNPs, ancestry- SNPs and whole mtDNA genomes are now available for casework use. Since RNA sequencing (RNA-Seq) is basically a form of targeted DNA sequencing (since it sequences targeted cDNAs that are created from mRNA) it should be a facile task to incorporate an RNA-Seq body fluid identification system into the NGS workflow. We reason that ‘democratizing’ personal genomics NGS technology might be a driver to provide forensic scientists with commercial solutions that will facilitate the routine capability of providing some activity level context to a DNA profile. Accordingly we have developed a targeted multiplexed next generation mRNA sequencing assay comprising a 33 biomarker panel that incorporates several highly specific gene targets with the necessary specificity to definitively identify most of the forensically relevant biological fluids.

## 2. Methods

### 2.1 Preparation of Body Fluid Stains

Overall, 232 samples were tested in the study (200 samples for development and performance checks and 32 for use in a blind study trial).

*Lab 1* - Body fluids were collected from volunteers using procedures approved by the University’s Institutional Review Board. Informed written consent was obtained from each donor. For development and performance testing, 152 samples were tested: blood (N=6), semen (N=6), saliva (N=6), vaginal secretions (N=6), menstrual blood (N =5), skin total RNA (N=3), two fluid mixtures (N=20), tissue total RNA (N=1 each of brain, lung, trachea, liver, skeletal muscle, heart, kidney, adipose, small intestine, stomach), sensitivity – 10, 5 and 1 ng inputs (two donors each of blood, semen, saliva, vaginal secretions and menstrual blood), reproducibility (N=30, two donors per body fluid each run in triplicate), species specificity (blood only; N=1 each of chimpanzee, baboon, mouse, duck, ferret, rabbit, guinea pig, cat, dog, deer, cow and pig), ‘non-standard’ samples (N=11, including menstrual blood samples collected over 6 days of reported menstruation, environmentally compromised samples and saliva swabbed from skin surface), DNA (N=4, 3 ng input) and three reaction blanks. Blood samples were obtained from commercial sources (Bioreclamation IVT (Long Island, NY), EDTA-containing vacutainers) and 50μl aliquots were dried onto cotton cloth. Freshly ejaculated liquid semen was provided in sealed plastic tubes and stored frozen until being dried onto sterile cotton swabs (IntegriSwabs, Lynn Peavey, Lenexa, KS) by placing full swab into liquid semen (i.e. saturation of cotton portion of the swab). Buccal samples (saliva) were collected from donors using sterile cotton swabs by swabbing the inside of the donor’s mouth. Semen-free vaginal secretions and menstrual blood were collected using sterile cotton swabs. Total RNA samples for skin (N=3) and tissues (N=1; brain, lung, trachea, liver, skeletal muscle, heart, kidney, adipose, small intestine and stomach) were purchased from commercial sources (ThermoFisher Scientific, Oyster Point, CA and BioChain^®^, Newark, CA). All samples were stored at -20°C or -80°C (commercially available tissue total RNA samples) until needed.

*Lab 2* – Body fluids were collected from healthy volunteers with their informed consent (N=48; eight samples each of blood, semen, saliva, vaginal secretions, menstrual blood and surface skin (7 from palm, one skin scraping)). The sampling was approved by the local ethics commission (KEK), declaration of no objection No. 24-2015. Blood was collected by finger prick, semen was collected in sterile cups and saliva was collected in sterile microtubes, and 50 μl was spotted onto a sterile cotton swab (Copan swabs, Millian AG, Nesselnbach/Niederwill, CH), respectively. Vaginal secretions and menstrual blood samples were retrieved from the vagina with sterile cotton swabs. Surface skin samples were collected by rubbing the skin surface with a pre-wetted (90% ethanol) sterile cotton swab. All swabs were dried at room temperature for at least 12 hours.

A set of 16 samples was analyzed by both laboratories and consisted of four blood (two of which were deposited on cellulose pads), two saliva (one of which was a buccal swab), two semen, one vaginal secretions, two menstrual blood and one skin sample (swab from palm of hand), as well as four two-fluid admixed samples: semen/vaginal secretions (1/4 vaginal swab with 12.5 μL semen), blood/saliva (50 μL each of blood and semen), semen/menstrual blood (1/4 menstrual blood swab with 12.5 μL semen) and saliva/skin (one skin swab with 50 μL saliva).

### 2.2 RNA Isolation

*Lab 1*- Total RNA was extracted from blood, semen, saliva, vaginal secretions and menstrual blood with guanidine isothiocyanate-phenol:chloroform (Ambion by ThermoFisher Scientific, Austin, TX) and precipitated with isopropanol [3]. Briefly, 500 μL of pre-heated (56°C for 10 min) denaturing solution (4M guanidine isothiocyanate, 0.02M sodium citrate, 0.5% sarkosyl, 0.1M β-mercaptoethanol) was added to a 1.5 mL Safe Lock extraction tube (Eppendorf, Westbury, NY) containing the stain or swab. The samples were incubated at 56°C for 30 min. The swab or stain pieces were then placed into a DNA IQ^TM^ spin basket (Promega, Madison, WI), re-inserted back into the original extraction tube, and centrifuged at 14,000 rpm (16,000 × g) for 5 min. After centrifugation, the basket with swab/stain pieces was discarded. To each extract the following was added: 50 μL 2 M sodium acetate and 600 μL acid phenol:chloroform (5:1), pH 4.5 (Ambion by ThermoFisher Scientific). The samples were then centrifuged for 20 min at 14,000 rpm (16,000 × g). The RNA-containing top aqueous layer was transferred to a new 1.5 mL microcentrifuge tube, to which 2 μL of GlycoBlue^TM^ glycogen carrier (ThermoFisher Scientific) and 500 μL of isopropanol were added. RNA was precipitated for 1 hour at −20°C. The extracts were then centrifuged at 14,000 rpm (16,000 × g) for 20 min. The supernatant was removed and the pellet was washed with 900 μL of 75% ethanol/25% DEPC-treated water. Following a centrifugation for 10 min at 14,000 rpm (16,000 × g), the supernatant was removed and the pellet dried using vacuum centrifugation (56°C) for 3 min. Twenty microliters of pre-heated (60°C for 5 min) nuclease free water (ThermoFisher Scientific) was added to each sample followed by an incubation at 60°C for 10 min. Extracts were used immediately or stored at -20°C until needed.

*Lab 2* – RNA was extracted with a guanidine isothiocyanate-phenol:chloroform extraction as described above [3]. A few samples (two to three of each body fluid type) were extracted using the QIAGEN RNeasy Mini kit or the Trizol reagent [26] according to manufacturer’s recommended protocol. To reduce the risk of RNase contaminations, RNase decontamination solution (RNaseZap^®^ RNase de-contamination solution, ThermoFisher Scientific, Zug, CH) and dedicated lab ware, pipettes and working space for RNA were used. Extracts were stored at -80°C until needed.

### 2.3 DNase I Digestion

*Lab 1* - DNase digestion was performed using the TURBO^TM^ DNA kit (ThermoFisher Scientific) according to the manufacturer’s protocol. Briefly, 1X TURBO^TM^ DNase Buffer and 1 μl TURBO DNase was added to the 20 μL RNA extracts and incubated at 37°C for 30 minutes and 75°C for 10 min (note: EDTA was not added during the heat activation step).

*Lab 2* – DNase digestion was performed using the TURBO DNA-free^TM^ kit (ThermoFisher Scientific) according to manufacturer’s recommended protocol (1 μL of DNase and 2.3 μl of inactivation buffer used).

### 2.4 RNA Quantitation

*Lab 1*- RNA extracts were quantitated with Quant-iT^TM^ RiboGreen^®^ RNA Kit (ThermoFisher Scientific) according to the manufacturer’s protocol. Fluorescence was determined using a Synergy^TM^ 2 Multi-Mode microplate reader (BioTek Instruments, Inc., Winooski, VT).

*Lab 2* - RNA extracts were quantitated with the QuantiFluor^®^ RNA System (Promega, Dubendorf, CH) on a Quantus^TM^ Fluorometer (Promega).

For both methods 2 μL of extract (20 μL, ~10%) was used for quantitation.

### 2.5 TruSeq^®^ Targeted RNA Library Preparation

NGS libraries of targeted body fluid gene candidates were prepared using the TruSeq^®^ Targeted RNA kit (January 2016 protocol version; Illumina Inc., San Diego, CA) and a TruSeq^®^ Targeted RNA custom oligonucleotide pool (referred to here as TOP) designed using Illumina Design Studio (see Table 1 for final 33-plex assay). All 48- or 96-sample thermal cycler reactions were performed on the Mastercycler^®^ pro S thermal cycler (Eppendorf, Hauppauge, NY) using thin-walled skirted Microseal^®^ PCR plates (BIO-RAD, Hercules, CA) sealed with Microseal^®^ B or A (for the amplification reaction) film (BIO-RAD). All 48- or 96-sample purification reactions (requiring the use of magnetic beads) were performed in 0.8 mL 96-well storage plates (ThermoFisher Scientific) and sealed with Microseal^®^ B film (BIO-RAD) and a magnetic stand- 96 (ThermoFisher Scientific). Lab 2 followed the same protocol, but used a GeneAmp^®^ PCR System 9700, MicroAmp^TM^ Optical 96-Well Reaction Plates, ABgene 1.2 mL 96-well storage plates and a MagJET 96-well Separation Rack (all ThermoFisher Scientific).

**Table 1.** Biomarker Composition of the 33-plex Targeted RNA NGS Multiplex for Body Fluid Identification.

| **Body Fluid** | **Gene Name** | **Assay ID** | **Chromosome** | **Transcript ID** | **Start Position** | **Stop Position** | **Product Size** |
| --- | --- | --- | --- | --- | --- | --- | --- |
| **Blood**<br>**(BL)** | **ALAS2** | 6678585 | X | NM_001037968 | 55046783 | 55044071 | 62 |
|  | **ANK1** | 6691928 | 8 | NM_001142446 | 41521209 | 41519420 | 73 |
|  | **SPTB** | 6758523 | 14 | NM_001024858 | 65289690 | 65271781 | 54 |
|  | **CD3G** | 6693072 | 11 | NM_000073 | 118220653 | 118221306 | 73 |
|  | **CD93** | 6973146 | 20 | NM_012072 | 23064962 | 23064906 | 58 |
|  | **AMICA1** | 6645104 | 11 | NM_001098526 | 118083145 | 118081400 | 51 |
| **Semen**<br>**(SE)** | **PRM1** | 6974413 | 16 | NM_002761 | 11374820 | 11374764 | 58 |
|  | **PRM2** | 6841914 | 16 | NM_002762 | 11369989 | 11369758 | 70 |
|  | **TGM4** | 6609462 | 3 | NM_003241 | 44935153 | 44938227 | 62 |
|  | **SEMG1** | 6747498 | 20 | NM_003007 | 43837335 | 43838280 | 76 |
|  | **SEMG2** | 6676715 | 20 | NM_003008 | 43852032 | 43852958 | 62 |
|  | **KLK3** | 6612360 | 19 | NM_001648 | 51358233 | 51359532 | 62 |
| **Saliva**<br>**(SA)** | **HTN3** | 6973356 | 4 | NM_000200 | 70902189 | 70902246 | 59 |
|  | **HTN1** | 6827099 | 4 | NM_002159 | 70921285 | 70924332 | 71 |
|  | **STATH** | 6820236 | 4 | NM_003154 | 70866669 | 70867937 | 63 |
|  | **PRB3** | 6653331 | 12 | NM_006249 | 11420141 | 11418925 | 71 |
|  | **PRB4** | 6810773 | 12 | NM_002723 | 11463303 | 11462311 | 64 |
|  | **PRH2** | 6683726 | 12 | NM_001110213 | 11081904 | 11082898 | 64 |
| **Vaginal**<br>**(VS)** | **CYP2B7P1** | 6700781 | 19 | NR_001278 | 41430314 | 41442027 | 61 |
|  | **DKK4** | 6702671 | 8 | NM_014420 | 42234489 | 42233315 | 71 |
|  | **FAM83D** | 6824170 | 20 | NM_030919 | 37570735 | 37576551 | 68 |
|  | **CYP2A6** | 6617506 | 19 | NM_000762 | 41352812 | 41351963 | 73 |
| **Menstrual**<br>**Blood**<br>**(MB)** | **MMP10** | 6711090 | 11 | NM_002425 | 102651246 | 102650443 | 63 |
|  | **LEFTY2** | 6721035 | 1 | NM_001172425 | 226127088 | 226125480 | 53 |
|  | **MMP7** | 6597729 | 11 | NM_002423 | 102394009 | 102391492 | 83 |
|  | **MMP11** | 6771272 | 22 | NM_005940 | 24122661 | 24122799 | 60 |
|  | **SFRP4** | 6784486 | 7 | NM_003014 | 37955727 | 37954015 | 74 |
| **Skin (SK)** | **LCE1C** | 6973810 | 1 | NM_178351 | 152777370 | 152777315 | 57 |
|  | **CCL27** | 6849211 | 9 | NM_006664 | 34662310 | 34662051 | 56 |
|  | **IL37** | 6656756 | 2 | NM_014439 | 113675327 | 113676166 | 57 |
|  | **SERPINA12** | 6730346 | 14 | NM_173850 | 94955989 | 94953798 | 67 |
|  | **KRT77** | 6768360 | 12 | NM_175078 | 53086239 | 53085733 | 60 |
|  | **COL17A1** | 6760475 | 10 | NM_000494 | 105840414 | 105838317 | 63 |

RNA was first transcribed into first strand cDNA following the TruSeq^®^ Targeted RNA kit degraded RNA protocol. The 10 μL reaction consisted of 7 μL of reaction mix (4 μL RCS1 buffer (Illumina Inc.), 2 μL ProtoScript^®^ II reverse transcriptase (New England Biolabs Inc., Ipswich, MA), 1 μL of 10X DTT (0.1M) (New England Biolabs)) and 3 μL of total RNA (target input approximately 50 – 100 ng, but varied slightly within each body fluid (averages for organic extraction: blood – 52 ng (N=10, 39 – 100 ng), semen – 47 ng (N=10, 14 – 123 ng), saliva – 67 ng (N=10, 15 – 129 ng), vaginal secretions – 81 ng (N=10, 50 – 106 ng), menstrual blood –86 ng (N=9, 50-126 ng) and surface skin – 18 ng (N=4, 11 – 26 ng)). The input (ng) quantities for the RNeasy Mini and Trizol extracted samples was generally lower than the target 50 ng total RNA input (averages: blood – 15 ng (N=4), semen – 19.7 ng (N=4), saliva – 10 ng (N=4) and surface skin – 17.6 ng (N=4)) with the exception of vaginal secretions and menstrual blood samples in which 50 ng of input total RNA was available. For sensitivity studies, total RNA inputs of 10, 5 and 1 ng were used (N=2 each for blood, semen, saliva, vaginal secretions and menstrual blood). Reaction plates were sealed and vortexed at 1600 rpm for 20 s and centrifuged at 280 × *g* for 1 min. Reverse transcription was performed as follows: 25°C 10 min, 42°C 30 min, 95°C 10 min and an infinite hold at 4°C.The cDNA samples were used immediately or stored at -20°C overnight (thawed at room temperature before subsequent use).

The custom TOP was next hybridized to the cDNA. The 10 μL hybridization reaction mix consisted of 5 μL TOP (Illumina Inc.) and 5 μL TE buffer pH 8.0 (ThermoFisher Scientific). Reaction plates were sealed and vortexed at 1600 rpm for 20 s. Following a 1-min incubation at room temperature, 30 μL of OB1 (Illumina Inc.) was added to each well. The plate was sealed and vortexed at 1600 rpm for 1 min. The 50 μL hybridization reactions were performed as follows: 70°C 5 min, 68°C 1 min, 65°C 2.5 min, 60°C 2.5min, 55°C 4 min, 50°C 4 min, 45°C 4 min, 40°C 4 min, 35°C 4 min, 30°C 4 min and a hold at 30°C. The bound oligos were then washed, extended and ligated according to the manufacturer’s protocol (TruSeq^®^ Targeted RNA, January 2016 protocol version; Illumina Inc.). The extension-ligation products were then amplified, adding Index 1 (i7) adapters and Index 2 (i5) adapters in the process. Each sample received a unique combination of i7 and i5 adapters in order to permit pooling of finished libraries prior to sequencing. Twenty microliters of the purified extension-ligation products were used in the 50 μL amplification reaction. The reaction plate was sealed, vortexed at 1600 rpm for 30 s and centrifuged at 280 x g for 1 min. The amplification reaction was performed as follows: 95°C 2 min; 34 cycles of 98°C 30 s, 62°C 30 s, 72°C 60 s; 72°C for 5 min and an infinite hold at 10°C. Amplification products were used immediately or stored at 4°C overnight if needed. The individual sample libraries were next purified according to the manufacturer’s protocol (TruSeq^®^ Targeted RNA, January 2016 protocol version; Illumina Inc.) resulting in a final sample library volume of 12.5 μL. Five microliters of each sample library were combined into a single pooled library per sequencing reaction. All individual libraries within a particular experiment (48 or 96 samples) were pooled into a single library. Pooled libraries and remaining individual libraries were stored at -20°C until needed.

### 2.6 TruSeq^®^ Targeted RNA Library Quantitation

Pooled libraries were quantitated using the 2200 TapeStation (Agilent Technologies, Santa Clara, CA) and High Sensitivity D1000 Screen tape according to the manufacturer’s protocol. Neat and 1:10 diluted libraries were run and the average concentration obtained from the 100-300 bp region was used to determine the library concentration (in nM).

### 2.7 MiSeq^®^ sequencing

Pooled libraries were diluted to 4 nM concentrations and denatured according to the manufacturer’s recommended protocol. Briefly, 5 μL of the 4 nM library was mixed with 5 μL 0.2 N NaOH and incubated at room temperature for 5 min. To the 10 μL denatured library sample, 990 μL of pre-chilled HT1 buffer (Illumina Inc.) was added resulting in a 20 pM sample. A 600 μL 6 pM sample was then prepared by further diluting the 20 pM library (180 μL 20 pM denatured sample and 420 μL pre-chilled HT1). The 600 μL 6 pM sample was immediately pipetted into the MiSeq^®^ v3 150 cycle reagent cartridge for sequencing on the MiSeq^®^ instrument (Illumina Inc.) using a v3 flow cell. The sequencing runs consisted of 51 single-end sequencing cycles. The average % Q30 scores for individual sequencing runs ranged from 95 – 98 percent, with % PF (passing filter) ranging from 85 – 96 percent and the % of PF reads identified is ~95%.

### 2.8 Data analysis

After sequencing, local sequencing software on the MiSeq analyzed the data (base calling, demultiplexing and alignment to the provided manifest file using a banded Smith Waterman algorithm) resulting in a target hits file that displays total reads per amplicon per sample. A minimum sample total read count (MTR) of 5,000 was used as an individual sample threshold and samples below the MTR were excluded from analysis. In addition a minimum biomarker read count (MBR) count of 500 was used as an individual biomarker threshold, with any counts below this threshold removed. A third threshold was then used in which individual biomarker read count values that were less than 0.5% of the total reads for the sample were also removed.

After filtering of samples in accordance with the above thresholds, the read count data was plotted in Microsoft^®^ Excel in order to create bar graphs of threshold-filtered counts by sample and by gene. The percent contribution of total reads (biomarker read count/total count for sample) was determined for each biomarker. The percent contribution of reads was next calculated in order to provide the percentage of total reads for each individual sample that was attributable to blood-, semen-, saliva-, vaginal secretions-, menstrual blood- and skin-biomarker specific markers and displayed as stacked bar graphs.

Agglomerative hierarchical clustering analysis is an alternative complementary method for data analysis that employs the raw hit counts as input without the use of ad hoc thresholds described above [27]. Clustering was performed using the BaseSpace^®^ TruSeq^®^ Targeted RNA v1.0 app (Illumina Inc.) which jointly clusters samples and biomarker amplicons. Briefly the software uses a minimum count threshold of 1, log-transforms the counts and performs median-normalization across all the samples. After clustering, the data are MAD-normalized so that the expression values for each gene are on the same scale. Biomarker amplicon and sample dendrogram files and a clustering heat map are used to visualize the similarities and differences in biomarker expression between samples.

## 3. Results

### 3.1 Assay Development

#### 3.1.1 Candidate Selection

Numerous putative tissue- or fluid-specific genes have been reported for most of the forensically relevant biological fluids [3,6–10,12,15,28–36]. While there is no definitive consensus set of genes required for use in forensic casework, there are some core genes for several body fluids that are routinely used (e.g. blood – ANK1, ALAS2, HBB; semen – PRM1, PRM2, TGM4, SEMG1; saliva – HTN3; vaginal secretions – CYP2B7P1; menstrual blood – MMP10, MMP7, LEFTY2) [5,8]. These genes, and others, have been extensively evaluated by the forensic community including numerous collaborative EDNAP studies on RNA profiling for body fluid identification [17,18,37–40], thus aiding our initial selection of genes for the targeted RNA sequencing assay. We designed and evaluated six targeted oligonucleotide primer pools (TOPs; TOP1 – 30-plex, TOP2 - 50-plex, TOP3 – 55-plex, TOP4 – 47-plex, TOP5 – 38-plex and TOP6 – 33-plex) which resulted in the testing of a total of 66 gene candidates for appropriate specificity (blood – 9 genes, semen – 7 genes, saliva – 18 genes, vaginal secretions – 10 genes, menstrual blood – 6 genes, skin – 13 genes and 3 housekeeping genes; data not shown).

Individual gene candidates were evaluated for specificity (e.g. ideal candidates with high read counts in target body fluid and low or no read counts in non-target body fluids) and abundance (e.g. ideal candidates with moderate to high read counts consistently amongst different donors of the target body fluid). Expression heat maps (Figure 1) were generated for each TOP design after initial testing to provide an easy visualization of gene expression between different body fluid sample types aiding in the selection of suitable candidates. Half of the initial gene candidates were deemed unsuitable based on factors such as poor performance (amplification efficiency), low abundance or cross-reactivity with non-target body fluids. Three housekeeping genes (B2M, SPRR3 and UBC [5,39]) were included in the first TOP design, only B2M was tested in TOP2 and subsequently TOP3 (and beyond) were devoid of housekeeping genes. The reason for excluding housekeeping genes was that they accounted for up to ~76% of the total reads for individual sequencing runs (e.g. for TOP1 ~ 78% of total reads out of ~10 million reads for the 24 body fluid samples tested were due to housekeeping genes (3.7 million reads for B2M and 4.1 million reads for SPRR3)) and their inclusion resulted in amplification and detection inefficiencies of the body fluid specific biomarkers. Similarly, extremely abundant biomarkers (such as HBB for blood) were also removed as they represented a large percentage of total reads making amplification of other biomarkers less efficient. Due to the high sensitivity of HBB, it was also being detected in a large number of samples where trace amounts of blood could be present (e.g. blood vessels, etc.).

**Figure 1.**
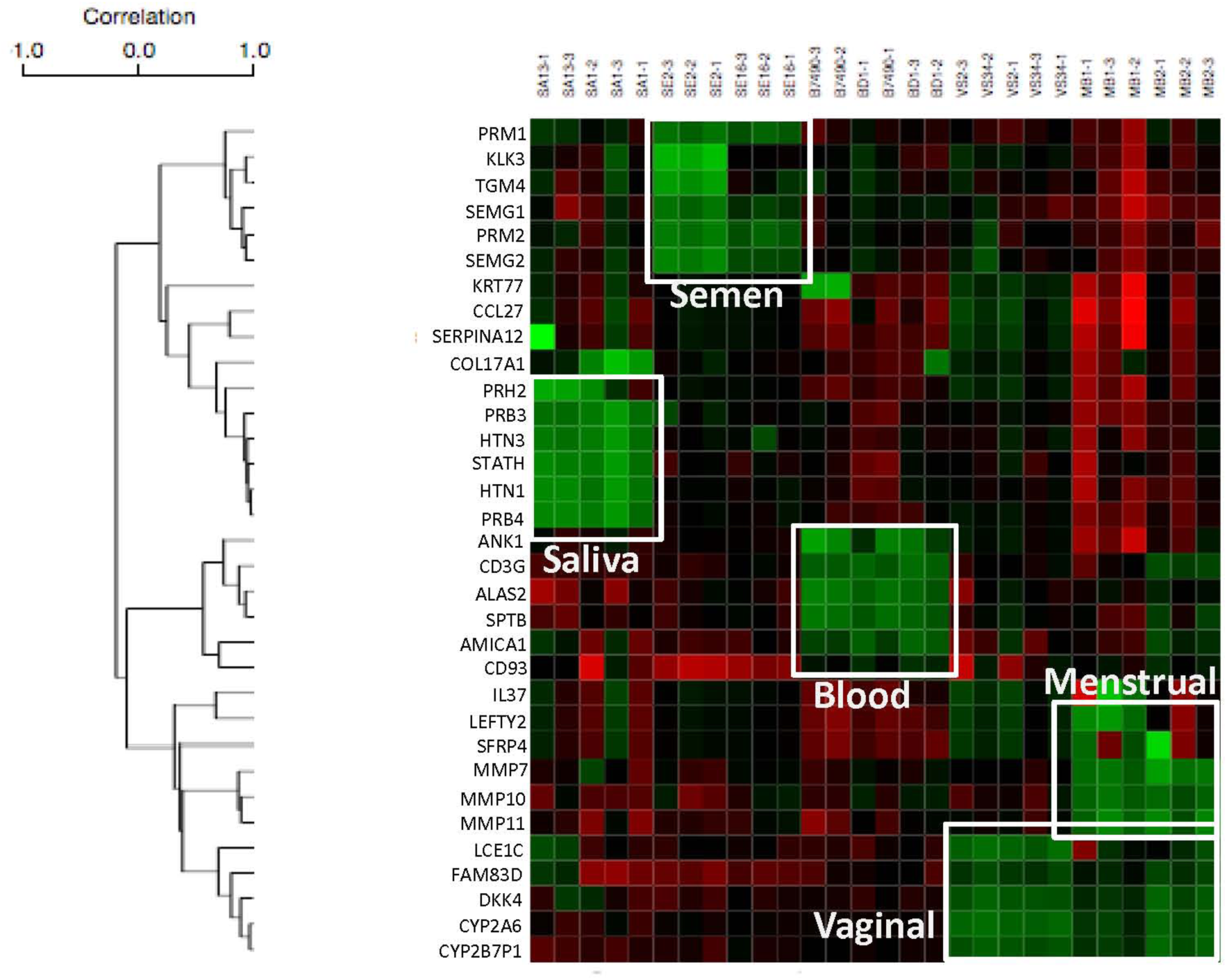
Gene Expression Heat map of 33 Body Fluid Specific Markers in Blood, Semen, Saliva, Vaginal Secretions and Menstrual Blood Samples. Y-axis – biomarker (gene) names. X-axis – body fluid samples (SA=saliva, SE=semen, B=blood, VS=vaginal secretions, MB=menstrual blood). Green represents higher expression, red represents lower expression. Clusters of up-regulated gene expression of a group of biomarkers specific to the target body fluid are highlighted with white boxes.

The above iterative selection process resulted in the development of a final 33-plex design (Table 1) that was deemed suitable for more extensive testing and evaluation that comprised six genes each for blood, semen, saliva and skin, five genes for menstrual blood and four for vaginal secretions.

#### 3.1.2 Specificity of the 33-plex Targeted RNA Sequencing Assay

The initial performance of the 33-plex targeted RNA sequencing assay was evaluated in total RNA samples prepared from a panel of 80 biological relevant fluids and tissues (blood, semen, saliva, vaginal secretions, menstrual blood, skin tissue and skin swabs. Input total RNA amounts were controlled but varied depending upon the sample; ~50 ng was determined to be most suitable. Raw read count data was evaluated and the following *ad hoc* thresholds employed: 1) minimum total read count (MTR) of 5,000 for individual samples, 2) minimum biomarker read count (MBR) of 500 as well as a 0.5% total read count threshold were used for individual biomarkers. Thus samples with total read counts below 5,000 were excluded from analysis. Read counts for individual biomarkers that were below 500 and that did not represent at least 0.5% of the total read counts for the sample were also removed (read count converted to 0). Six of the 80 samples were excluded based on these thresholds: one blood, two semen, one saliva and two skin swab samples.

The read count values for each biomarker were averaged amongst the 74 samples in order to evaluate the specificity of the included biomarkers (Table 2). Amongst the 13 <u>blood</u> donors tested, the average read count for blood biomarkers ranged from 95,469 (ALAS2) to 17,945 (CD3G). Expression was not observed for a majority of the non-blood biomarkers, with detectable counts only observed for PRM2 (2,756 average counts in 2 of 13 donors), SEMG1 (3,353 average counts in 2 of 13 donors) and FAM83D (4,690 average count from 3 of the 13 donors).

**Table 2.** Body Fluid Specificity of 33 gene candidates. Average read counts of each biomarker in body fluids and skin (calculated from N donors). For each body fluid sample set the average input (ng) and average total read counts are listed. Number in parentheses represents the number of samples in which the biomarker was detected. Shading: dark grey ≥ 10,000 read counts; light grey 5,001 – 9,999 read counts; no color≤5,000 read counts.

|  | Body Fluid | Blood | Semen | Saliva | Vaginal | Menstrual | Skin tissue | Skin touch |
| --- | --- | --- | --- | --- | --- | --- | --- | --- |
|  | *N* | *13* | *12\** | *13* | *14* | *13* | *3* | *6* |
|  | *Avg ng input* | 40.4 | 39.9 | 53.2 | 72.0 | 74.9 | 3.0 | 19.4 |
|  | *Avg total* | 280,635 | 1,313,629 | 149,164 | 526,188 | 803,671 | 60,228 | 34,575 |
| BL | ALAS2 | 95,469<sub>(13)</sub> |  |  |  | 4,589<sub>(8)</sub> |  |  |
|  | ANK1 | 44,133<sub>(12)</sub> |  |  |  | 3,952<sub>(4)</sub> |  |  |
|  | SPTB | 24,813<sub>(13)</sub> |  |  |  | 1,572<sub>(2)</sub> |  |  |
|  | CD3G | 17,945<sub>(11)</sub> |  |  |  | 3,718<sub>(4)</sub> |  |  |
|  | CD93 | 34,757<sub>(12)</sub> |  | 9,949<sub>(7)</sub> | 16,861<sub>(8)</sub> | 14,784<sub>(12)</sub> |  |  |
|  | AMICA1 | 61,714<sub>(13)</sub> |  | 8,487<sub>(7)</sub> | 12,773<sub>(8)</sub> | 4,645<sub>(7)</sub> |  | 964<sub>(2)</sub> |
| SE | PRM1 |  | 844,872<sub>(11)</sub> | 1,785<sub>(3)</sub> | 1,715<sub>(2)</sub> | 1,272<sub>(4)</sub> |  | 3,089<sub>(3)</sub> |
|  | PRM2 | 2,756<sub>(2)</sub> | 396,666<sub>(11)</sub> |  |  |  |  | 1,369<sub>(3)</sub> |
|  | TGM4 |  | 58,194<sub>(10)</sub> |  |  |  |  |  |
|  | SEMG1 | 3,353<sub>(2)</sub> | 155,505<sub>(10)</sub> |  |  |  |  |  |
|  | SEMG2 |  | 69,212<sub>(8)</sub> |  |  |  |  |  |
|  | KLK3 |  | 4,740<sub>(5)</sub> |  |  |  |  |  |
| SA | HTN3 |  |  | 110,657<sub>(13)</sub> | 4,951<sub>(2)</sub> |  |  | 4,879<sub>(2)</sub> |
|  | HTN1 |  |  | 6,316<sub>(9)</sub> |  |  |  |  |
|  | STATH |  |  | 10,632<sub>(11)</sub> |  |  |  |  |
|  | PRB3 |  |  | 3,823<sub>(8)</sub> |  |  |  |  |
|  | PRB4 |  |  | 5,723<sub>(5)</sub> |  |  |  |  |
|  | PRH2 |  |  | 2,197<sub>(5)</sub> |  |  |  |  |
| VS | CYP2B7P1 |  |  | 4,375<sub>(6)</sub> | 185,176<sub>(14)</sub> | 139,934<sub>(11)</sub> | 1,259<sub>(2)</sub> | 804<sub>(2)</sub> |
|  | DKK4 |  |  | 1,165<sub>(4)</sub> | 28,774<sub>(13)</sub> | 37,116<sub>(9)</sub> |  |  |
|  | FAM83D | 4,690<sub>(3)</sub> |  | 7,345<sub>(8)</sub> | 231,361<sub>(14)</sub> | 179,006<sub>(12)</sub> | 1,170<sub>(2)</sub> |  |
|  | CYP2A6 |  |  |  | 36,120<sub>(11)</sub> | 25,983<sub>(10)</sub> |  |  |
| MB | MMP10 |  |  | 2,146<sub>(3)</sub> | 4,084<sub>(4)</sub> | 296,332<sub>(12)</sub> |  | 3,175<sub>(3)</sub> |
|  | LEFTY2 |  |  |  |  | 34,332<sub>(5)</sub> |  |  |
|  | MMP7 |  |  | 3,358<sub>(2)</sub> | 12,443<sub>(3)</sub> | 196,963<sub>(7)</sub> |  |  |
|  | MMP11 |  |  |  |  | 7,393<sub>(4)</sub> |  |  |
|  | SFRP4 |  |  |  |  |  |  |  |
| SK | LCE1C |  |  | 1,057<sub>(2)</sub> | 43,278<sub>(11)</sub> | 22,566<sub>(7)</sub> | 25,961<sub>(3)</sub> | 22,684<sub>(6)</sub> |
|  | CCL27 |  |  |  |  |  | 12,012<sub>(3)</sub> |  |
|  | IL37 |  |  |  |  |  | 4,593<sub>(2)</sub> | 5,544<sub>(2)</sub> |
|  | SERPINA12 |  |  |  |  |  | 3,684<sub>(2)</sub> | 5,951<sub>(3)</sub> |
|  | KRT77 |  |  |  |  | 790<sub>(3)</sub> | 3,079<sub>(3)</sub> |  |
|  | COL17A1 |  |  |  |  |  | 9,479<sub>(3)</sub> |  |

\*1/12 semen samples from a vasectomized male; PRM1/PRM2 therefore only expected in 11/12 samples.

For <u>semen</u>, detectable expression was only observed for the included semen biomarkers, with average count values ranging from 844,872 (PRM1) to 58,194 (TGM4). The average count value for KLK3 (4,740) was significantly lower than the other semen biomarkers and was only detected in 5 of the 12 donors.

For <u>saliva</u>, average counts were overall much lower than the biomarkers for the other body fluids. HTN3 was expressed the highest with an average count of 110,657 and was detected for all 13 donors. There was low level expression observed for nine non-saliva biomarkers with the most significant being CD93 (average count of 9,949 amongst 7 of the 13 donors), AMICA1 (average count of 8,487 amongst 7 of the 13 donors) and FAM83D (average count of 7,345 amongst 8 of the 13 donors). It is interesting to note that expression of CD93 and AMICA1 were only observed in data from one of the two participating laboratories and all data generated was from a single run. These count values are significantly lower than the HTN3 average count.

For <u>vaginal secretions</u>, high average read counts were obtained for all four vaginal secretions biomarkers amongst the 14 donors tested with the highest expression observed for CYP2B7P1 (average read count of 185,176 amongst all 14 donors) and FAM83D (average read count of 231,361 amongst all 14 donors). Expression of the skin biomarker LCE1C was observed in 11 of the 14 vaginal secretions samples. This is frequently observed in vaginal samples and is likely due to the genuine presence of skin in these samples as they are self-collected by donors not wearing gloves and contact with external skin surfaces is possible during the self-collection process. Higher than expected read counts were observed for CD93 (average read count of 16,861 amongst 8 of 14 donors), AMICA1 (average read count of 12,773 amongst 8 of the 14 donors) and MMP7 (average read count of 12,443 amongst only 3 of the 14 donors).

For <u>menstrual blood</u>, four of the five biomarkers were detected, with no read counts above thresholds for SFRP4. All four vaginal secretions biomarkers and all six blood biomarkers were also expressed in the menstrual blood samples as would be expected since menstrual blood is comprised of varying ratios of menstrual blood, peripheral blood and vaginal secretions.

For the <u>skin</u> total RNA samples (i.e. skin tissue total RNA purchased commercially), the highest expression was observed for LCE1C (average read count of 25,961amongst all 3 donors), CCL27 average read count of 12,012 amongst all three donors) and COL17A1 (average read count of 9,479 amongst all three donors). For <u>surface skin</u> samples, LCE1C had the highest average read count (22,684 amongst all six donors) with additional expression of only IL37 and SERPINA12 amongst the 6 skin biomarkers. The slightly differing expression profiles for skin total RNA samples and surface skin samples is expected as the biomarkers were selected for detection specifically for both surface skin/touch samples (e.g. LCE1C) and skin tissue (e.g. CCL27) realizing that all six biomarkers may not be detected in each skin sample type.

In addition to analyzing the raw read count values, we also plotted the threshold-filtered read counts in several simple bar graph formats for easy visualization of the differential gene expression of the biomarkers. Supplementary Figure 1 shows expression data graphed ‘by gene’ (one representative gene selected for each of the target body fluids) for a sub-set of the body fluid samples (29 of the 74 body fluids samples; sub-set of samples used only for easier viewing of the sample types on the x-axis). As can be seen from the expression data, the biomarkers are present in the target body fluid with little to no cross-reactivity with non-target body fluids. It is interesting to note that for PRM2 (Supplementary Figure 1B), 0 counts are observed for the first semen donor. This is semen from a vasectomized male and therefore was not expected to show expression of the sperm cell specific marker PRM2. Figure 2 shows 33-gene expression data for individual body fluid samples. As can be seen from these graphs, highly specific gene expression profiles were observed with a majority of expression originating from the biomarkers deemed specific to that fluid or tissue. The biomarker and sample-type graphs indicate the high degree of specificity of the prototype assay.

**Figure 2.**
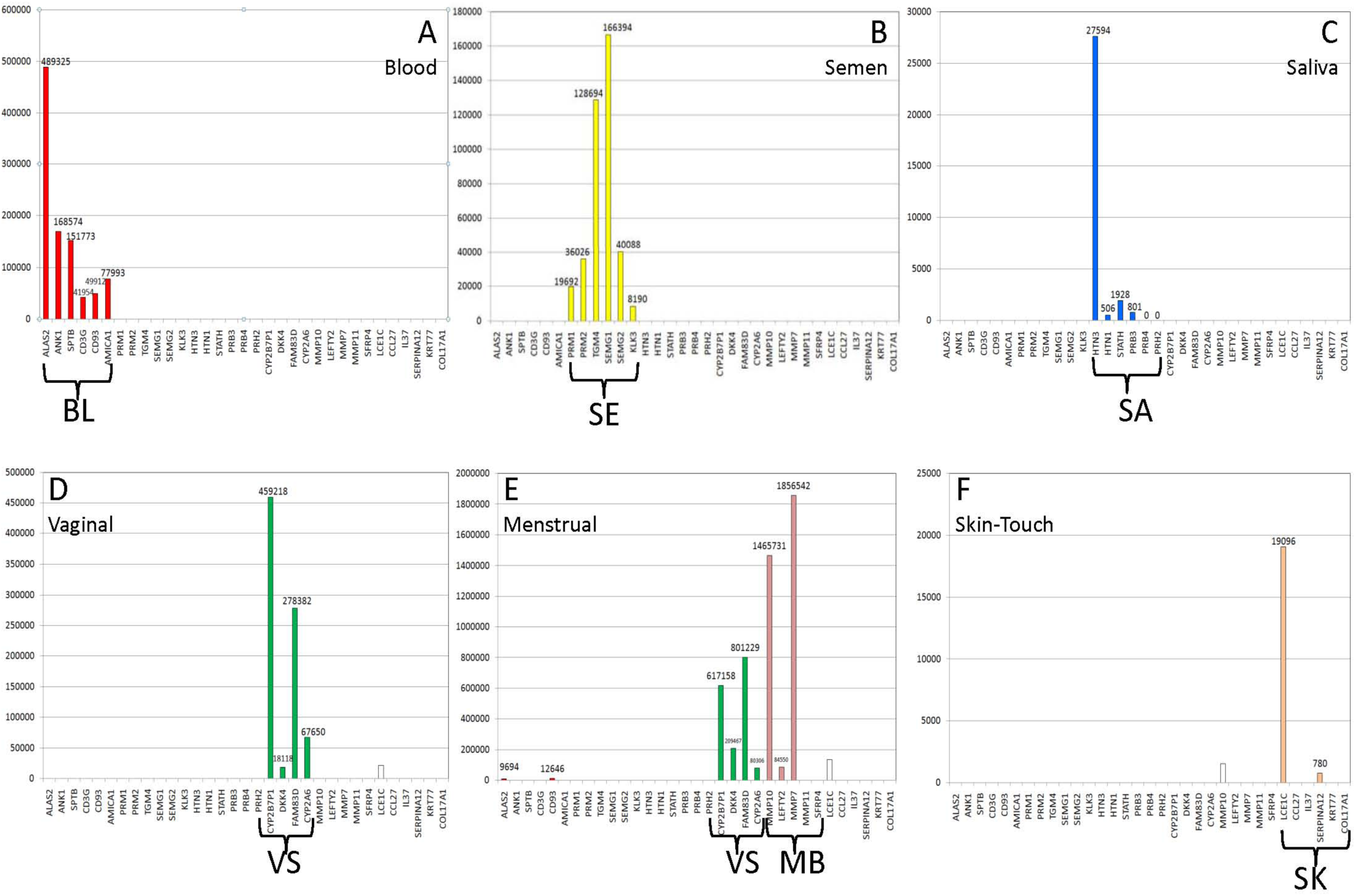
Gene Expression Profiles for Different Individual Body Fluid Types Using the 33-plex Targeted RNA Sequencing Assay. Read counts for 33 body fluid specific genes are shown for individual body fluid samples (A – blood, B – semen, C – saliva, D – vaginal secretions, E – menstrual blood, F – skin). Colored bars represent expression of body fluid specific biomarkers within the target body fluid (red – blood, yellow – semen, blue – saliva, green – vaginal secretions, pink – menstrual blood, peach – skin). For full reference to colors, readers are directed to the online version of the article. White bars represent expression of a biomarker not classified as specific to that body fluid. Y-axis – read counts, X-axis – body fluid specific genes.

#### 3.1.3 Body Fluid Inference

Inference of the presence of a body fluid based upon quantitative gene expression (as used in our targeted RNA sequencing assay) is a classification problem whose goal is to determine a body fluid category for an unknown sample. Here we have investigated the use of a couple of *ad hoc* binary approaches to body fluid prediction whose output is a simple categorical statement of the presence (or absence) of a particular body fluid. Specifically these include (i) assigning the percentage of reads in a sample that are due to each of the 6 tissues/body fluids categories tested and (ii) inter-sample differential gene expression revealed by unbiased agglomerative hierarchical clustering.

Filtered read counts for individual biomarkers were divided by the total reads for the sample and then expressed as a percentage. The percentages from each of the biomarkers deemed specific for an individual fluid were then added together. The percent composition of the total reads attributable to each class of biomarkers is then visualized as bar graphs. The percent biomarker class composition values for a sub-set of 12 samples (two donors for each body fluid) are shown in Figure 3. As can be seen from this figure, 100% of the total reads for the blood samples are attributable to biomarkers of the blood class. The same is observed for semen with 100% of the total reads for the semen samples attributable to the semen class of biomarkers. While Figure 3 shows only a sub-set of samples analyzed using this approach, the same process was repeated for all 74 samples included in the analysis. The average percent contributions for each class of biomarkers were determined for each body fluid (Table 3). For blood, on average 93% of total reads for blood samples was attributable to blood class biomarkers. For semen, 100% of the total reads for semen samples were attributable to semen class biomarkers. For menstrual blood, menstrual blood class biomarkers accounted for 49% of the total reads for menstrual blood samples, with an additional 43% attributed to vaginal class biomarkers and 7% from blood class biomarkers. For vaginal secretions, on average 85% of total reads were attributable to vaginal secretions class biomarkers with 7% attributable to skin (likely for reasons described previously). Seventy-one to eighty-five percent of total reads for saliva and skin (both skin swab and skin tissue samples) were attributable to the biomarker classes specific to that body fluid or tissue type.

**Figure 3.**
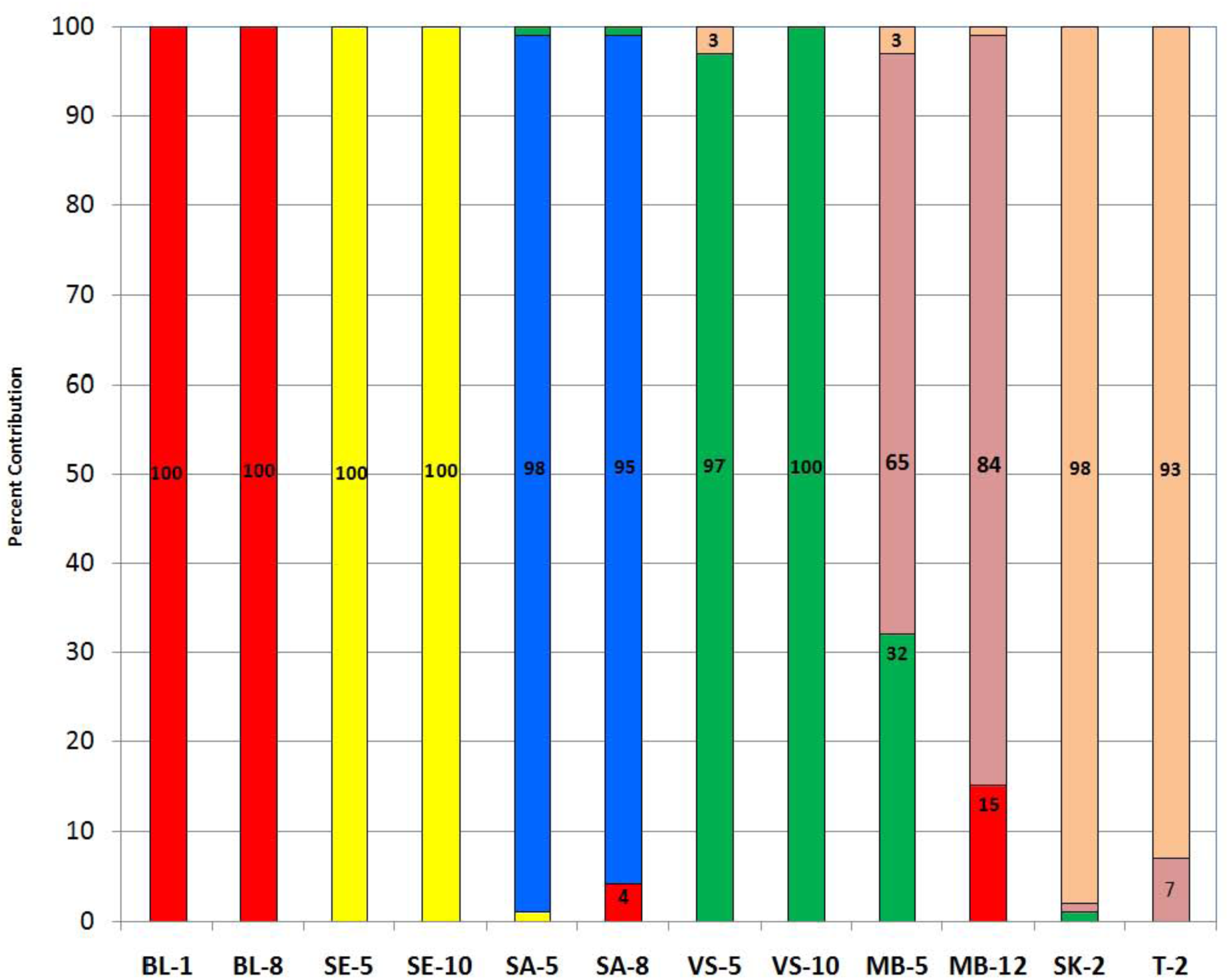
Biomarker Expression Composition for Individual Body Fluid Samples. The percent contribution to the sample for each individual biomarker was calculated (reads per biomarker/total reads per sample). The percentages from each body fluid specific biomarker were combined into pre-determined classes according to their adjudged specificity in order to determine the percentage of reads per sample attributable to each body fluid or tissue class. For example, 100% indicates that all reads for a sample were attributable to that class of body fluid specific biomarkers. Percent reads attributable to each biomarker class are listed and represented by color: red – total expression from the six biomarkers comprising the blood class, yellow – total expression from the six semen biomarkers comprising the semen class, blue – total expression from the six saliva biomarkers comprising the saliva class, green – total expression from the 4 vaginal secretions biomarkers comprising the vaginal secretions class, pink – total expression from the 5 menstrual blood biomarkers comprising the menstrual blood class and peach – total expression from the 6 skin biomarkers comprising the skin class. Y-axis – percent contribution; X-axis – body fluid samples (BL=blood (N=2), SE=semen (N=2), SA=saliva (N=2), VS=vaginal secretions (N=2), MB=menstrual blood (N=2), SK or T=skin (N=2)).

**Table 3.** Contribution of Body Fluid Biomarker Classes to Different Body Fluids. Average percent contributions of each biomarker class is shown for each body fluid (BL – blood biomarker class comprises 6 different individual gene markers; SE – semen biomarker class comprises 6 different individual gene markers; SA – saliva biomarker class comprises 6 different individual gene markers; VS – vaginal secretions biomarker class comprises 4 different individual gene markers; MB – menstrual blood biomarker class comprises 5 different individual gene markers; SK – skin biomarker class comprises 6 different individual gene markers). The number of donors (N) used to determine the averages are provided.

|  | Blood | Semen | Saliva | Vaginal | Menstrual | Skin | Touch |
| --- | --- | --- | --- | --- | --- | --- | --- |
| Biomarkers | *N=13* | *N=12* | *N=13* | *N=14* | *N=13* | *N=3* | *N=6* |
| BL | **93** | 0 | 10 | 4 | **7** | 2 | 1 |
| SE | 3 | **100** | 1 | 1 | 0 | 2 | 9 |
| SA | 3 | 0 | **81** | 0 | 0 | 0 | 8 |
| VS | 1 | 0 | 6 | **85** | **43** | 10 | 1 |
| MB | 0 | 0 | 2 | 3 | **49** | 1 | 9 |
| SK | 0 | 0 | 0 | 7 | 1 | **85** | **71** |
| Total | 100 | 100 | 100 | 100 | 100 | 100 | 100 |

Using this approach of apportioning the relative amounts of the different classes of biomarkers in a sample, the gene expression data is essentially normalized and is not dependent upon the total read counts, which will vary from sample to sample. Sample total reads will vary for a variety of reasons but principally from different input amounts into the assay (50 ng of total RNA is not always possible to input since only 3 μl of sample extract can be used in the assay at present and different samples may not yield 50 ng in 3 μl). Additionally using this approach, it would not be necessary for all biomarkers specific to a specific body fluid or tissue to be expressed in order for an accurate source identification to be made. This is important as expression of some of the lower abundance biomarkers are not obtained for every sample of the specific body fluid.

Evaluation of the similarities and differences in gene expression of the 33 targeted genes between samples was performed using agglomerative hierarchical clustering analysis. The clustering was performed jointly on samples and biomarker amplicons using unfiltered raw read counts. Results indicate that the 33-gene assay exhibits a high degree of specificity for body classes in that samples of the same body fluid type cluster together due to similarities in gene expression. Figure 4 shows a representative dendrogram of the unbiased clustering of body fluid types when samples from skin (n=2), saliva (n=5), semen (n =6), blood (n=5), menstrual blood (n =5) and vaginal secretions (n=5) were analyzed. As can be seen from the clusters the six different body fluid/tissue classes show clear and distinct intra-class differences in gene expression, whereas samples of the same body fluid /tissue type cluster together. Interestingly the vasectomized male shown in the dendrogram clusters with semen samples as expected.

**Figure 4.**
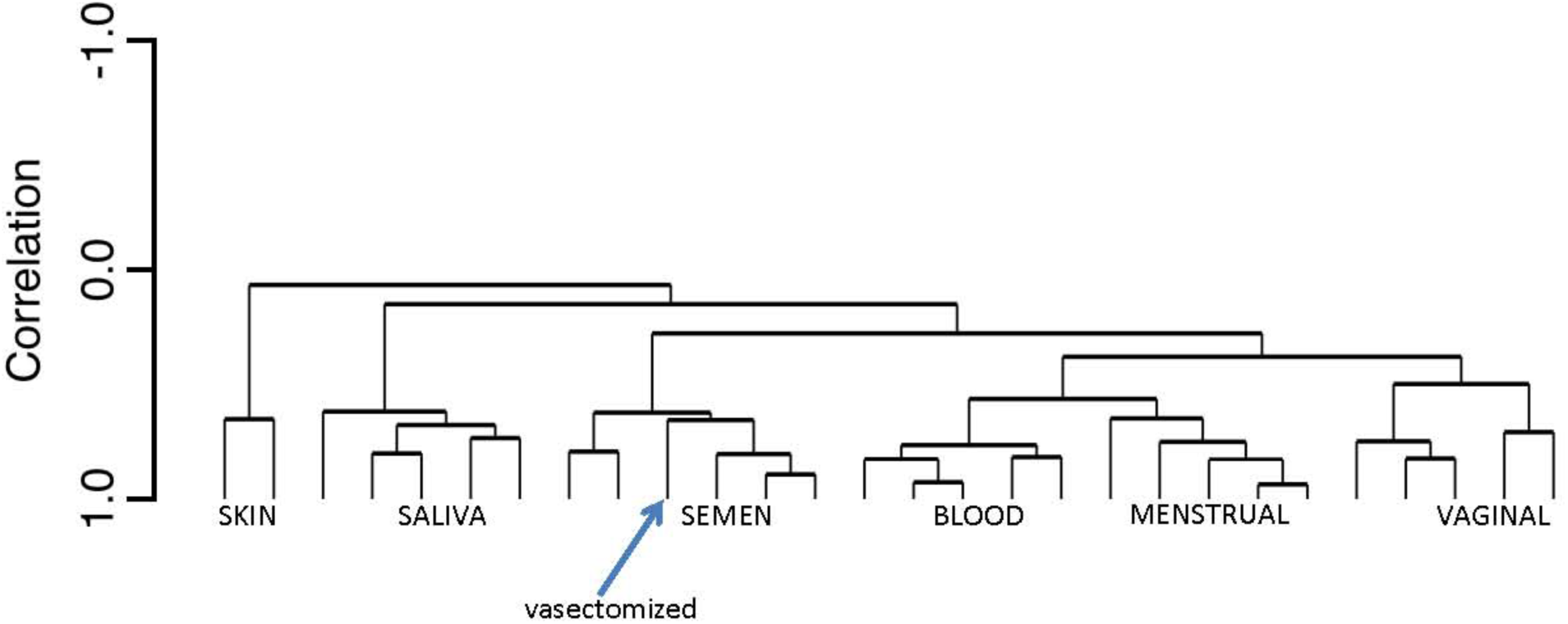
Dendogram of Single Source Samples Clustering According to Similarities in Gene Expression. The arrow indicates the position of a semen sample from a vasectomized male. The gene expression correlation distance between samples is indicated by the length of the vertical branch points on the Y axis.

### 3.2 Performance testing

We have carried out an initial set of performance checks on the prototype targeted RNA assay to ascertain its efficacy for potential use in forensic casework.

#### 3.2.1 Biomarker input sensitivity

The current target input amount of total RNA for the assay is 50 ng, which is the same as the manufacturer’s recommended quantity. To evaluate the differing sensitivity of individual biomarkers to detection, we analyzed 10, 5 and 1 ng total RNA input quantities for two donors each of blood, semen, saliva, vaginal secretions and menstrual blood. The results of the sensitivity study are summarized in Supplementary Table 1. Saliva and vaginal secretions biomarkers were undetectable at 10 ng or below. For blood, ALAS2 was readily detectable with both 10 and 5 ng input total RNA amounts. Expression was also detectable for four additional blood biomarkers (ANK1, SPTB, CD3G and AMICA1) although average read counts were low (~1,000 – 4,000). For semen, PRM1 and PRM2 were readily detectable using 10 and 5 ng input total RNA which is indicative of the high abundance of these two biomarkers in sperm. TGM4 was also detectable with ~4,000 – 5,000 reads using 5 and 10 ng of input total RNA, respectively. Low but detectable reads were also obtained for SEMG1 and KLK3 for both inputs. For menstrual blood, read counts of ~3,000 – 5,000 were obtained for MMP10, CYP2B7P1 and FAM83D using 10 ng of input total RNA. No results were obtained for 1 ng input of total RNA for any of the body fluids.

These results provide an indication of the current sensitivity levels of the included biomarkers, but do not necessarily provide an accurate estimate of the limit of detection (LOD) of the assay. The use of NGS for RNA profiling is still in early development and, as with any new technique, it is expected that sensitivity will improve as additional optimization work is performed. Thus 50 ng remains a reasonable target input amount for the current prototype assay although some, but not all, body fluids were detectable with 5-10 ng total RNA input. Numerous other samples throughout the study also contained less than the optimal 50 ng and successful results were obtained.

#### 3.2.2 Mixtures

During the commission of a violent crime two or more people are present and may have the opportunity of depositing biological material at the crime scene or onto an individual or individuals, thus resulting in a mixture of biological material (i.e. body fluids and/or tissues). Comprehensive body fluid identification assays developed for forensic use must have the ability to detect multiple body fluids and tissues in admixed samples. We performed some preliminary testing of the targeted RNA assay on a limited number of possible binary fluid admixture samples including blood/saliva, vaginal secretions/semen, semen/saliva and blood/semen. Three separate experiments were conducted.

In the first experiment two different body fluid mixture types (blood/saliva and vaginal secretions/semen) were prepared such that in each sample there was a constant amount of the first body fluid with a varying proportion of the second. Two sets of samples per mixture type were prepared that comprised body fluids from different pairs of donors. The blood/saliva samples were prepared by adding liquid saliva (10, 5 and 1 μl aliquots) to ½ portions of 50 μl dried blood stains on cotton and the vaginal secretions/semen samples were prepared by adding liquid semen (10, 5 and 1 μl aliquots) to ¼ portions of dried vaginal swabs respectively. The determined percent body fluid class composition for each of these mixtures types after targeted RNA analysis is shown in Figure 5A.

**Figure 5.**
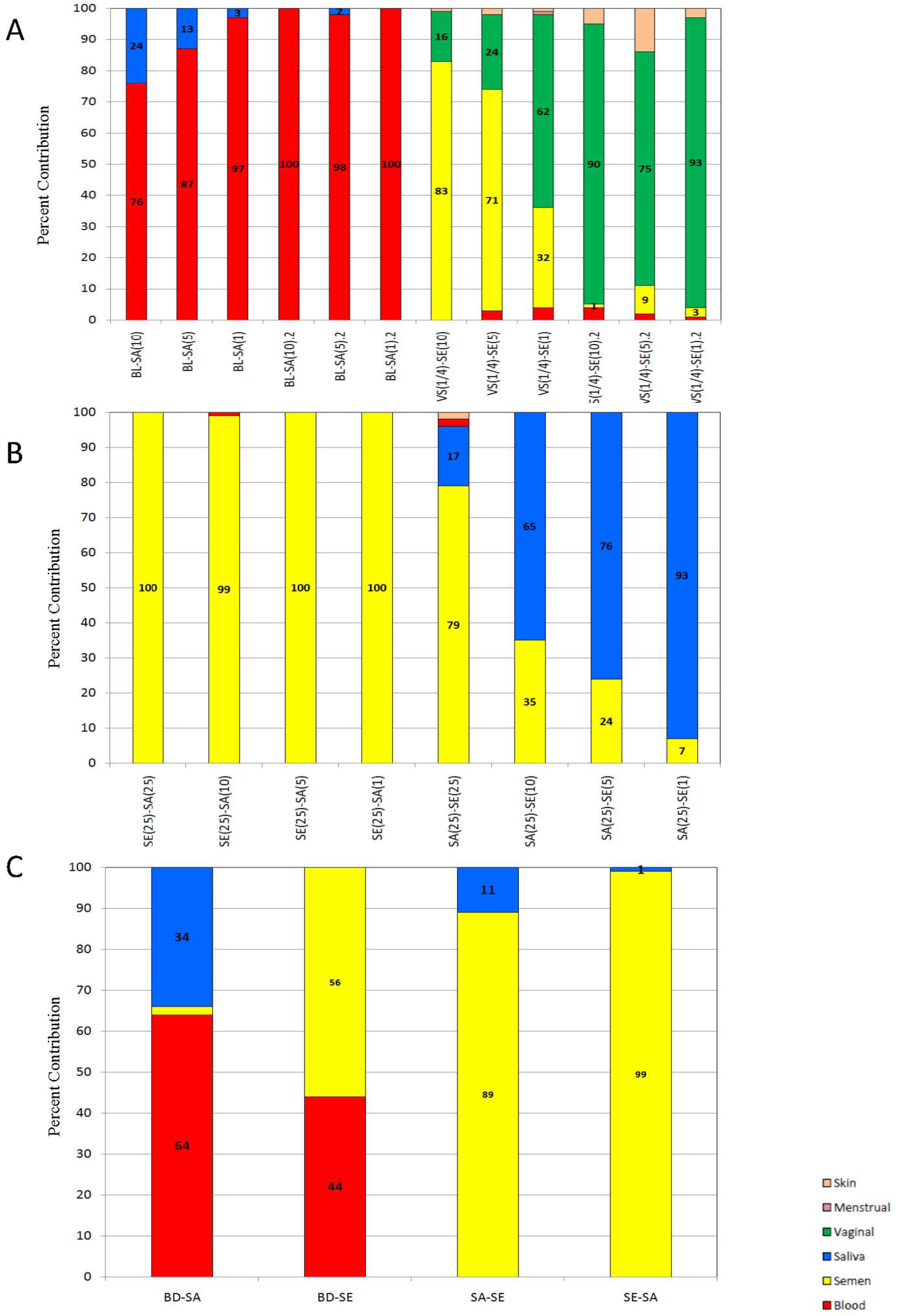
Identified Biomarker Expression Classes in Two-Fluid Admixed Body Fluid Samples. The percent contribution for individual biomarkers was calculated (reads per biomarker/total reads per sample). The percentages from each group of body fluid specific biomarkers were combined in order to determine the percentage of reads per sample attributable to each body fluid or tissue class. Percent reads attributable to each biomarker group are listed and represented by color: red – expression from blood biomarkers, yellow – expression from semen biomarkers, blue – expression from saliva biomarkers, green – expression from vaginal secretions biomarkers, pink – expression from menstrual blood biomarkers and peach – expression from skin biomarkers. A) blood-saliva and vaginal-semen admixed samples (two donor sets for each mixture type); liquid saliva (10, 5 and 1 μl aliquots) added to ½ of a 50 μl dried blood stains and liquid semen (10, 5 and 1 μl aliquots) added to ¼ dried vaginal swabs. B) semen-saliva and saliva-semen admixed samples (one donor set for each mixture type); liquid body fluids combined as 25-25 μl, 25-10 μl, 25-5 μl and 25-1 μl volumes (volume of each body fluid listed in parentheses in each sample name). C) blood-saliva (BD-SA) - 50 μl liquid saliva added to a 50 μl dried blood stain; blood-semen (BD-SE) – 10 μl liquid blood mixed with 1 μl liquid semen; saliva-semen (SA-SE)– 15 μl liquid saliva mixed with 5 μl liquid semen; semen-saliva (SE-SA) – 15 μl liquid semen mixed with 5 μl liquid saliva. All mixtures were dried overnight prior to analysis. Y-axis – percent contribution; X-axis – body fluid samples (BL=blood, SE=semen, SA=saliva, VS=vaginal secretions, MB=menstrual blood, SK=skin).

For the first donor set in the blood/saliva mixtures, blood biomarkers represented 76, 87 and 97% of the total reads for the mixture samples in which decreasing quantities of liquid saliva was added. Saliva biomarkers represented the remaining 24, 13 and 3% of total reads, respectively. Thus this sample set exhibited the expected pattern of gene expression with only blood and saliva biomarker classes detected, and with the proportion of the sample comprising the saliva biomarker class decreasing in accord with the decreased proportion of liquid saliva that had been added to the mixture. However, for the second blood/saliva donor set, saliva was essentially undetected with the percent of total reads from blood biomarkers accounting for 98 – 100% of the total reads. For the mixture in which 5 μl of liquid saliva was added, a small (2%) contribution was attributable to saliva class biomarkers, a level that probably would be indistinguishable from background sequence noise.

The results for the vaginal secretions/semen mixtures illustrate the transcriptional heterogeneity that can arise due to the presence of vaginal secretions in mixtures. In the first donor set, expression of semen class biomarkers represented 83, 71 and 32% of the total reads for the samples in which decreasing quantities of liquid semen had been added to the vaginal swabs. The expression of vaginal biomarkers in these samples concomitantly increased as the ratio of semen to vaginal secretions decreased, with 16, 24 and 62% of the total reads attributable to vaginal secretions biomarkers. Similar to the blood/saliva mixtures, for donor set two of the vaginal secretions/semen, the minor component (i.e. semen) was barely detected with the semen biomarkers only comprising 1, 9 and 3 % of the total reads present in which 10, 5 and 1 μl of liquid semen, respectively, was added. In addition to the expected vaginal secretions and semen biomarkers, all of the samples expressed skin biomarkers (1-15%) and most (5/6) expressed low levels (5%) of blood biomarkers. The low level expression of skin and blood biomarkers as well as high expression of vaginal secretions biomarkers in vaginal secretions samples is a characteristic of this body fluid (Table 2) and will have to be taken into account when inferring body fluid types in a mixture.

In the second mixture experiment semen/saliva mixtures were tested in which liquids from both body fluids were mixed at different ratios, dried and then subjected to targeted RNA analysis. The two body fluids were combined as follows: 25-25 μl, 25-10 μl, 25-5 μl and 25-1 μl volumes. In one set (designated ‘semen/saliva’), semen at a constant amount (25 μl) was diluted with different saliva volumes (25-1 μl). In the other set (designated ‘saliva/semen’) saliva was present at a constant 25 μl while semen was added in decreasing volumes (25-1 μl). For the semen/saliva mixtures in which semen was the major component, 99 – 100% of the total reads were attributable to semen biomarkers (Figure 5B). The semen biomarkers are highly expressed and are therefore relatively sensitive compared to the saliva biomarkers (Supplementary Table 1) and there is a significant difference in the amount of RNA in equal volumes of semen and saliva (more total RNA per unit volume in semen). Therefore it was not entirely unexpected that semen biomarker expression could consume the read counts in this mixture. For the saliva/semen mixtures, however, in which saliva was the major component, both semen and saliva were identified (Figure 5B). For the samples in which 25, 10, 5 and 1μl of liquid semen was added, the percent contribution from saliva biomarkers was 17, 65, 76 and 93%, respectively. The respective percent contribution from semen biomarkers was 79, 35, 24 and 7%. This mixture set clearly showed the expected decrease in semen contribution with a corresponding increase in saliva biomarker contribution as the volume of semen added to the saliva was decreased.

In the third experiment four separate mixtures with known added amounts of body fluids were analyzed: blood/saliva in which 50 μl liquid saliva was added to a 50 μl dried blood stain, blood/semen in which 10 μl liquid blood was mixed with 1 μl liquid semen, saliva/semen (3:1) in which 15 μl liquid saliva was mixed with 5 μl liquid semen, and semen/saliva (3:1) in which 15 μl liquid semen was mixed with 5 μl liquid saliva. The results are shown in Figure 5C. For the blood/saliva mixture, both body fluids were successfully detected with 64 and 34% contributions of the total reads, respectively. Two percent of the reads were attributable to semen biomarkers and are likely due to background analytical or transcriptional noise. For the blood/semen mixture, both body fluids were also successfully detected with 44 and 56% contributions of the total reads due to blood and semen, respectively. For the saliva/semen (3:1) mixture, the presence of both body fluids was identified with 11 and 89% of the total reads attributable to biomarkers for saliva and semen, respectively. A smaller contribution of the total reads was observed for saliva in this mixture even though by volume it is the major component. Despite the 3:1 saliva/semen ratio, due to the significantly higher amounts of RNA recovered from semen samples as well as the high abundance of the semen biomarkers, semen expression still represented a greater percentage of the total reads. However saliva was still detectable at a level sufficiently above background. For the semen/saliva (3:1) mixture in which semen was the major component, 99% of the total reads were attributable to semen biomarkers with only 1% attributable to saliva. This level of expression is indistinguishable from background noise and the absence of robust expression of saliva in some semen/saliva mixtures is consistent with the observations made in experiment two above.

#### 3.2.3 Repeatability

Single source body fluid samples (two donors for each body fluid) were run in triplicate in order to assess the reproducibility of the developed assay. Library preparation and sequencing for all triplicates was performed at the same time in order to avoid any potential run-to-run variation. For each triplicate set, we examined total reads per sample, read counts for each individual biomarker and percent contributions to the total reads from individual biomarkers and from the body fluid biomarker class (Figure 6 and Supplementary Table 2 and Table 3).

**Figure 6.**
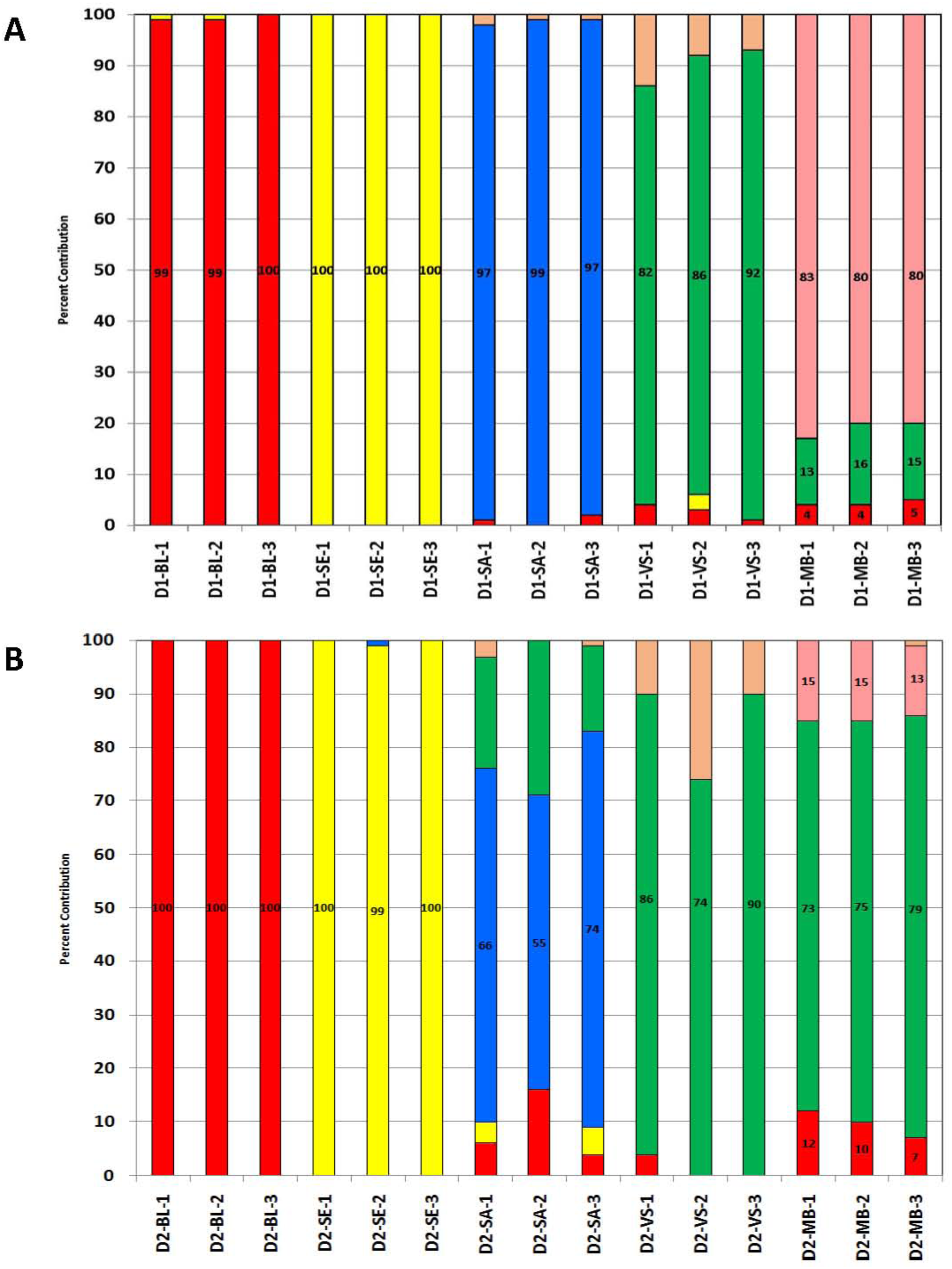
Repeatability of the NGS Targeted RNA Assay in Triplicate Sampling of Individual Body Fluid Samples. Individual body fluid samples were run in triplicate (-1, -2, -3; A) Donor 1 (D1); B) Donor 2 (D2)) using the same amount of input (in ng). The percent contribution for individual biomarkers was calculated (reads per biomarker/total reads per sample). The percentages from each group of body fluid specific biomarkers were combined in order to determine the percentage of reads per sample attributable to each body fluid or tissue class. Percent reads attributable to each biomarker group are listed and represented by color: red – expression from blood biomarkers, yellow – expression from semen biomarkers, blue – expression from saliva biomarkers, green – expression from vaginal secretions biomarkers, pink – expression from menstrual blood biomarkers and peach – expression from skin biomarkers. Y-axis – percent contribution; X-axis – body fluid samples (BL=blood, SE=semen, SA=saliva, VS=vaginal secretions, MB=menstrual blood, SK=skin).

Significant variation was observed for the read count data between replicates (Supplementary Table 3). The percent composition analysis normalizes these values to the total read counts and as expected greater consistency between replicates was observed. Figure 6 provides a graphical representation of percent contributions of the biomarkers in each body fluid class for each triplicate sample from both donors. The data clearly shows good repeatability of the targeted RNA assay across the triplicate samples. For example, with blood and semen 99 – 100% of biomarker expression in all three replicates was attributable to blood and semen biomarker classes, respectively. In general, vaginal secretions replicates had the highest coefficient of variation (CV) (6 and 10% for donor 1 (D1) and 2 (D2) respectively), with the exception of one of the two saliva samples (SA-D2) in which the CV for percent composition of target biomarkers was 15% (Supplementary Table 2). The saliva sample highlights the expected observation that greater variation between replicates will occur for those samples where amplification efficiency is reduced (i.e. those samples with lower overall total reads) For SA-D2, the total percent contribution in donor 2 from saliva biomarkers ranged from 55 to 74% amongst the three replicates in contrast to the SA-D1 sample from donor 1 in which the total percent contribution ranged from 97-99% with a CV of only 1%. The vaginal sample in which a CV of 10% was observed (VS-D2) also had very low overall total reads for each of the replicates (average of only ~11,000 total reads).

In summary, the assay repeatability was good when the raw counts were normalized as a percent of the total reads and apportioned into the appropriate body fluid specific classes.

#### 3.2.4 ‘Non-standard’ samples

A small number of ‘non-standard’ samples were analyzed in order to further test the performance of the targeted RNA sequencing assay. These samples included an evaluation of menstrual blood biomarker expression over a 6-day menstruation period, environmentally compromised samples and, as a mock casework sample, a saliva stain swabbed from the surface of an arm.

The percent contributions of biomarkers for the relevant body fluid classes of biomarkers (menstrual blood, vaginal secretions and blood) were determined for menstrual blood samples collected daily through days 1 – 6 of a self-reported menstruation period from a single donor (Supplementary Figure 2). Expression of menstrual blood biomarkers was very high on days 1 and 2 (95 and 93% of total read counts) and then was significantly reduced on days 3 – 6, down to 1 and 0% of total reads by days 5 and 6, respectively. Vaginal secretions biomarkers were low on days 1 – 5 but then increased by day 6, which corresponds to the time at which menstrual blood biomarkers disappeared. Peripheral blood biomarkers only accounted for 5 – 24% of the total reads across the 6 day period, with the highest expression (24%) observed on day 4. Since only one donor set was available for testing, definitive general conclusions cannot be made regarding expression of these biomarkers in other individuals or indeed in the same individual. However, the variation in observed biomarker expression is in agreement with expectations based upon the well-characterized physiological events during menses.

Several environmentally compromised samples were tested to provide an initial assessment of assay performance with challenging and possibly degraded samples (Supplementary Figure 3). Separate bloodstains from the same donor (i) stored in the back seat of a car for 4 days and (ii) incubated at 30°C with 90% humidity for 1 week were analyzed and blood was successfully detected in both samples (percent contribution of total reads attributable to the blood biomarker class was 100 and 99%, respectively). The presence of semen was confirmed (100% read contribution from semen biomarkers) for a known semen stain on denim that was incubated at 37°C for 3 months. Vaginal secretions biomarkers were identified in a vaginal swab sample incubated at 37°C for 1 year (75% read contribution from vaginal secretions biomarkers).

A saliva/skin mixture was also tested in which saliva was applied to the surface of an arm by licking and allowed to dry (Supplementary Figure 3). This type of sample may be encountered in sexual or physical assault cases. All of the biomarker expression in this sample (100%) was attributable to saliva biomarkers.

These initial results from body fluid stains kept at elevated temperatures and over long periods of time and from skin deposits are promising with respect to the potential utility of the targeted RNA for use with challenging or compromised samples in casework.

#### 3.2.5 Species specificity

Blood samples were available for the following primates and animals: chimpanzee, baboon, mouse, rabbit, guinea pig, cat, duck, ferret, dog, deer, cow and pig (N=1 for each). An average input of ~43 ng total RNA was used for testing the targeted RNA assay with different non-human species samples. Dried bloodstain samples from duck, ferret, dog, deer, cow and pig were tested with none of these samples resulting in total read counts above the 5,000 minimum sample count threshold. The read counts for blood biomarkers from chimpanzee, baboon, mouse, rabbit, guinea pig and cat blood samples are provided in Supplementary Table 4. ANK1 and CD93 were not detected in any of the primate or animal samples. Very low counts were observed for AMICA1 for chimpanzee and cat (1,987 and 571, respectively), in comparison to ~61,000 read counts for human blood samples. ALAS2 was detected in chimpanzee (13,252 reads), baboon (788,672 reads) and rabbit (867,005 reads) samples. SPTB was also highly expressed in mouse (15,542 reads), rabbit (58,703 reads) and cat (11,037 reads). CD3G was also highly expressed in baboon (96,782 reads). Expression in the chimpanzee and baboon samples was expected, as these are higher primates. Amongst the above threshold animal samples, the only significant animal cross-reactivity observed was for SPTB (detected in mouse, rabbit, cat) and ALAS2 in rabbit. However, none of the other blood biomarkers were present in these animal species. Therefore, while the expression of SPTB and ALAS2 does not appear to be exclusive to humans and primates, it may be possible to use the overall pattern of expression of blood biomarkers to differentiate humans and primates although this would require testing additional samples of each primate or animal species and performing differential gene expression analysis. Non-human species testing of the other (non-blood) biomarkers was not possible due to difficulties in obtaining a sufficient number of samples of the other body fluids from animals.

**Table 4.**
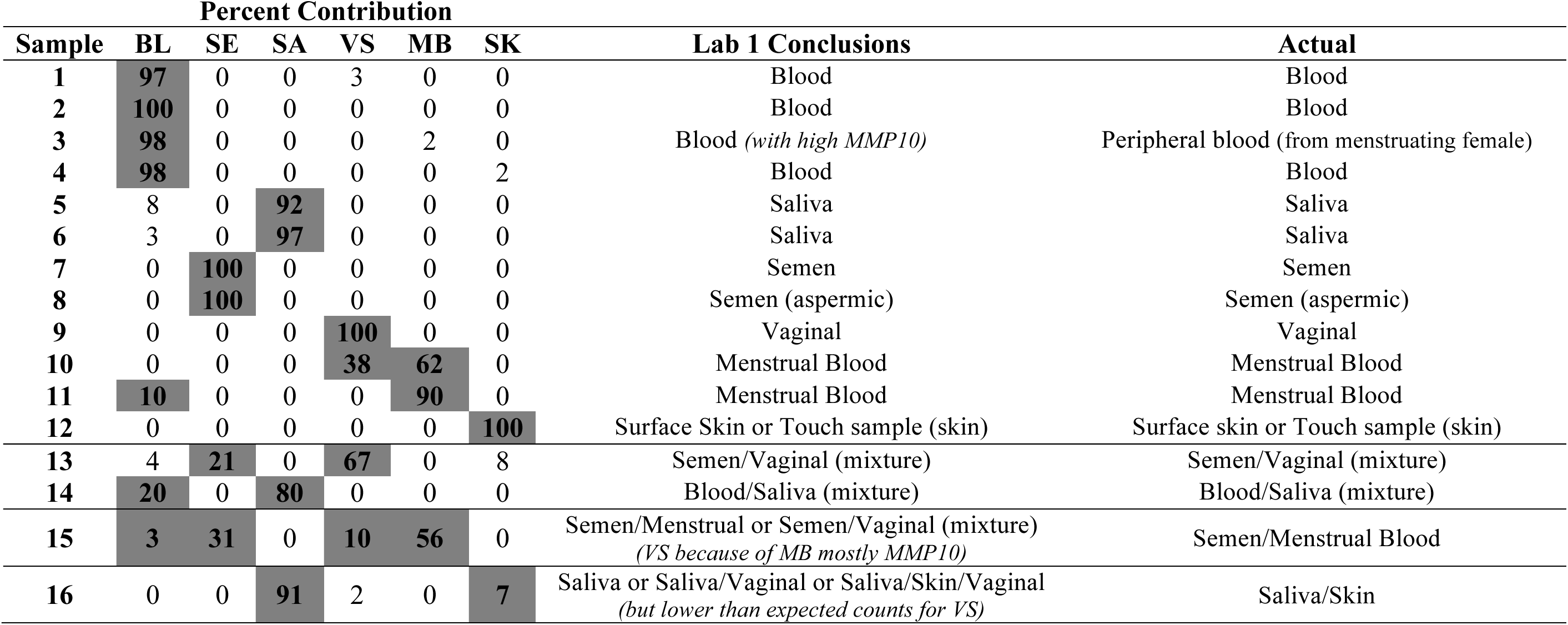
Body Fluid Identification in a 16-Sample Blind Study Using the 33-plex Targeted RNA Sequencing Multiplex.

| Sample | Percent Contribution |  |  |  |  |  | Lab 1 Conclusions | Actual |
| --- | --- | --- | --- | --- | --- | --- | --- | --- |
|  | BL | SE | SA | VS | MB | SK |  |  |
| 1 | 97 | 0 | 0 | 3 | 0 | 0 | Blood | Blood |
| 2 | 100 | 0 | 0 | 0 | 0 | 0 | Blood | Blood |
| 3 | 98 | 0 | 0 | 0 | 2 | 0 | Blood (*with high MMP10*) | Peripheral blood (from menstruating female) |
| 4 | 98 | 0 | 0 | 0 | 0 | 2 | Blood | Blood |
| 5 | 8 | 0 | 92 | 0 | 0 | 0 | Saliva | Saliva |
| 6 | 3 | 0 | 97 | 0 | 0 | 0 | Saliva | Saliva |
| 7 | 0 | 100 | 0 | 0 | 0 | 0 | Semen | Semen |
| 8 | 0 | 100 | 0 | 0 | 0 | 0 | Semen (aspermic) | Semen (aspermic) |
| 9 | 0 | 0 | 0 | 100 | 0 | 0 | Vaginal | Vaginal |
| 10 | 0 | 0 | 0 | 38 | 62 | 0 | Menstrual Blood | Menstrual Blood |
| 11 | 10 | 0 | 0 | 0 | 90 | 0 | Menstrual Blood | Menstrual Blood |
| 12 | 0 | 0 | 0 | 0 | 0 | 100 | Surface Skin or Touch sample (skin) | Surface skin or Touch sample (skin) |
| 13 | 4 | 21 | 0 | 67 | 0 | 8 | Semen/Vaginal (mixture) | Semen/Vaginal (mixture) |
| 14 | 20 | 0 | 80 | 0 | 0 | 0 | Blood/Saliva (mixture) | Blood/Saliva (mixture) |
| 15 | 3 | 31 | 0 | 10 | 56 | 0 | Semen/Menstrual or Semen/Vaginal (mixture)<br>(*VS because of MB mostly MMP10*) | Semen/Menstrual Blood |
| 16 | 0 | 0 | 91 | 2 | 0 | 7 | Saliva or Saliva/Vaginal or Saliva/Skin/Vaginal<br>(*but lower than expected counts for VS*) | Saliva/Skin |

#### 3.2.6 Organ Tissue specificity

In addition to the body fluid samples, the expression of each of the included biomarkers was evaluated using a panel of 10 human tissues (Supplementary Table 5). Since the tissue total RNA samples were obtained from commercial sources and of extremely high purity, an input of 1 ng total RNA was used. Total reads for lung, heart, kidney, adipose and stomach did not meet the minimum count (MTR) threshold. The read counts for brain, small intestine, trachea, liver and skeletal muscle are provided in Supplementary Table 5. Significant expression was only detected for four biomarkers: ANK1, SPTB, PRB3 and PRB4. ANK1 was detected in brain and SPTB in brain and skeletal muscle. The detection of blood in these samples is likely *bona fide* detection of trace amounts of blood remaining in the tissue samples from blood vessels after tissue dissection and RNA isolation (e.g. blood vessels, etc). PRB3 and PRB4 were detected in trachea samples. As these are saliva biomarkers, it is possible that some saliva biomarkers could be found in an anatomically connected tissue such as trachea due to drainage from the mouth at post mortem. However the saliva biomarker HTN3 was not detected in trachea and this might be able to be used to distinguish saliva from trachea if PRB 3 and PRB 4, after further more detailed studies, are found to be truly expressed in trachea. Thus saliva would express HTN3 and often PRB 3 and PRB4, although trachea would express PRB 3 and PRB 4 in the absence of HTN3.

#### 3.2.7 DNA and Amplification Blanks

The RNA sample preparation process includes a DNase treatment step in order to remove any residual DNA that may be present in the RNA extracts. However, it is possible that even with the DNase treatment a small amount of residual DNA may be present. Therefore, genomic DNA itself was tested with the targeted RNA assay to determine whether any products from contaminating DNA (e.g. from processed pseudo-genes) would be produced that could confound RNA biomarker analysis and interpretation. An input amount of 3 ng was selected since it is not anticipated that significant amounts of genomic DNA would survive DNase treatment. The DNA samples were carried through the full targeted RNA sequencing process, including reverse transcription.

Four DNA samples were tested. An average sample read count of 505 was observed for the DNA samples, with read counts for individual samples ranging from 0 to 2,761 reads. None of the DNA samples had total read counts above the minimum 5,000 read count threshold and therefore would be excluded from analysis due to it being considered noise.

Three amplification blanks were included in the targeted RNA sequencing experiments performed. The amplification blanks contained water in place of any sample. None of the amplification blanks had total read counts above the minimum 5,000 read count (two had 0 counts and one had a 1,514 read count for MMP10) and therefore would be excluded from any analysis.

### 3.3 Blind Study

A set of 16 samples was prepared by Lab 2 and sent to Lab 1 to analyze as a blind study. Lab 1 had no knowledge of the sample type prior to conducting their analysis. Lab 2 tested the same samples separately as well to evaluate reproducibility of the assay between the two laboratories. The read count data obtained for the set of 16 samples from both laboratories is shown in Supplementary Table 6. The raw read count data was filtered using the MTR and MBR thresholds and the percent contributions from each biomarker body fluid class were determined for each sample (Figure 7, Table 4). The conclusions made by Lab 1 are provided in Table 4.

**Figure 7.**
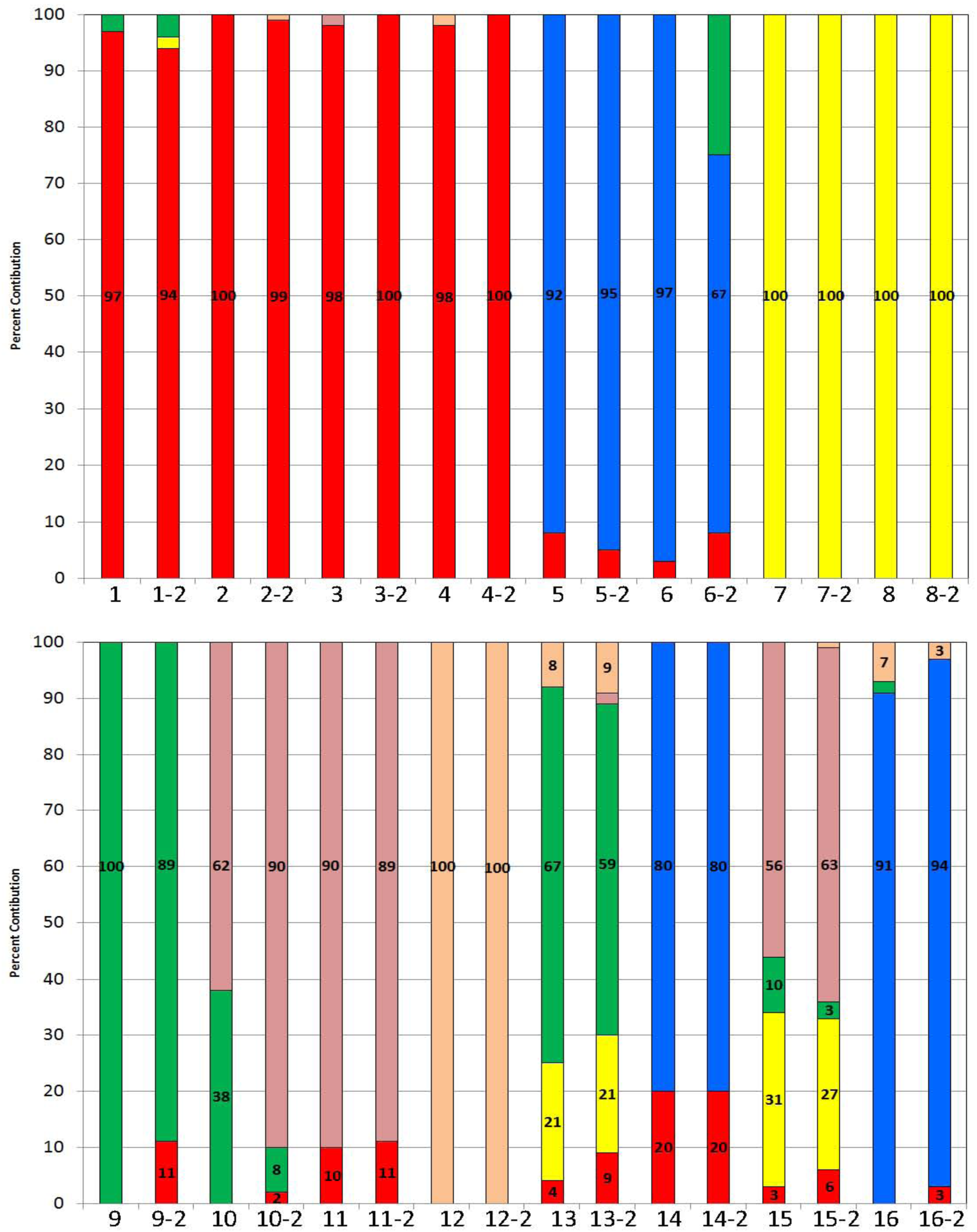
Identification of 16 Unknown Body Fluid Samples using the 33-plex Targeted RNA Sequencing Assay. Sixteen body fluid samples of known provenance were prepared by Lab 2 and submitted to Lab 1 as a blind study in which the nature of the samples was unknown to Lab 1. Samples were numbered 1-16. The ‘reference’ Lab 2 also tested the same set of samples (results designated with ‘-2’ labels in the sample names). For each sample, the percentage of reads per sample attributable to each body fluid or tissue class was determined. Percent reads attributable to each biomarker class are listed and represented by color: red – expression from blood biomarkers, yellow – expression from semen biomarkers, blue – expression from saliva biomarkers, green – expression from vaginal secretions biomarkers, pink – expression from menstrual blood biomarkers and peach – expression from skin biomarkers. Y-axis – percent contribution; X-axis – sample identifier.

Samples 1 – 12 appeared to be single source body fluids samples. Samples 1 – 4 were identified as blood since the contribution from blood biomarkers was 97 – 100% for each of those samples. Sample 3 had a small percentage of reads attributable to menstrual blood biomarkers. However, since it was only 2% of the total reads (and was solely from MMP10) a menstrual blood component was not inferred. Samples 5 and 6 were identified as saliva with 92 and 97% of total reads attributable to saliva biomarkers. Samples 7 and 8 were identified as semen, with sample 8 identified as semen originating from an aspermic donor as 0 counts were observed for the sperm cell markers PRM1 and PRM2 (Supplementary Table 6). Sample 9 was identified as vaginal secretions, with 100% of the total reads attributable to vaginal secretions biomarkers. Samples 10 and 11 were identified as menstrual blood as they possessed 62 and 90% of total reads attributable to menstrual blood biomarkers. Sample 10 also showed the presence of vaginal secretions and sample 11 showed the presence of blood, which is also consistent with the presence of menstrual blood in these samples. Sample 12 had 100% of total reads attributable to skin biomarkers and was therefore identified as a skin sample. LCE1C was the only biomarker detected for this sample and therefore this sample was inferred as likely being a surface skin or touch sample. As can be seen from the actual results in Table 4, the single source body fluid inferences made by Lab 1 were all correct. Interestingly, sample 3 which had a higher MMP10 read count (although only for Lab 1) was peripheral blood that originated from a menstruating female.

Samples 13 and 14 were identified as a semen/vaginal secretions mixture (21% semen biomarkers, 67% vaginal biomarkers) and a blood/saliva mixture (20% blood biomarkers, 80% saliva biomarkers), respectively. Sample 15 was also identified as an admixed sample. Lab 1 identified this sample as either semen/menstrual blood or semen/vaginal secretions. The contribution from menstrual blood biomarkers in the menstrual blood class was almost all from MMP10, which is sometimes elevated in vaginal samples [12]. Blood biomarkers accounted for 3% of the total reads but since this is a low percentage it was not a factor in the interpretation. With the actual source of the sample being menstrual blood, the contribution from blood biomarkers was genuine. Review of Lab 2 results for this sample included significant menstrual blood read counts for MMP10, MMP7 and SFRP4, which would be more characteristic for the presence of menstrual blood compared to just vaginal secretions. Sample 16 was a mixture, although Lab 1 conclusions couldn’t exclude the possibility of saliva only, a saliva/vaginal secretions mixture or a saliva/skin/vaginal secretions mixture. The actual sample was a saliva/skin mixture. The presence of saliva was definitive with 91% of the total reads attributable to saliva biomarkers. However, the presence of vaginal secretions or skin or both could not be excluded. Skin biomarkers accounted for 7% of the total reads. However, an additional 2% were from vaginal biomarkers. A higher skin percentage is often observed for vaginal samples and therefore it was included as a possible result. This sample illustrates the need for analytical thresholds to be established for the identification of minor component fluids in admixed samples.

Since both laboratories completed the analysis of these 16 samples, we were able to evaluate the reproducibility of the assay. This was an insightful comparison since different laboratories with slightly different protocols and different analysts completed the work. Figure 7 shows a side-by-side comparison of the biomarker class compositions for each sample from both laboratories (‘-2’ samples from Lab 2). As can be seen, there is a high degree of reproducibility in the data generated by both labs. In fact, there were only two samples, 9 and 10, where there were some more noticeable differences (>10%) in the minor fluid contributions although the main body fluid component was still very clear and consistent with the known composition for both samples.

## 4. Discussion

We report the development of a prototype next generation sequencing (NGS) targeted mRNA profiling assay for body fluid identification designed to definitively identify 5 body fluids (blood, semen, saliva, menstrual blood and vaginal secretions) and skin. The 33 biomarkers present in the assay were chosen after iterative specificity testing of a number of candidate gene transcripts. Each biomarker is grouped into one of six different body fluid/tissue classes with each class comprising 4-6 specific markers. Subsequent to initial specificity testing during assay design, a number of other performance checks were carried out on the prototype assay to assess its accuracy and precision. The assay is demonstrated here to be highly specific with minimal or no confounding cross reactivity of the biomarkers with non-targeted body fluids. In terms of throughput the assay takes two days to perform manually without liquid handling robots and we routinely process 48 samples at the same time although we have on occasion successfully tested 96. 384 wells are available and, according to the manufacturer, suitable for the targeted RNA assay but liquid handling robots would be preferable to use to process this number of samples. We believe the current prototype assay shows promise in being ‘fit for purpose’ but recognize that further optimization, testing and evaluation needs to be performed before it should be used in forensic casework. The use of liquid handling robots for pre-sequencing sample and library preparation would significantly reduce hands on time and improve assay repeatability and reproducibility.

Although the targeted RNA assay is based upon the selection of up-regulated biomarkers in specific body fluids, intra-class variation in expression of biomarkers within each class is observed. Some biomarkers exhibit high expression as exemplified by ALAS2 (blood), PRM1 (semen), HTN3 (saliva), CYP2B7P1 (vaginal secretions), MMP10 (menstrual blood) and LCE1C (skin). Others have lower expression (often ten times lower or more than the highest expresser) such as CD3G (blood), KLK3 (semen), PRB3 (saliva), DKK4 (vaginal secretions), LEFTY2 (menstrual blood) and IL37 (skin). The presence of high and low expressers might be advantageous to have in that the presence of both the high and low expressers in a sample might indicate that the input quantity and quality of sample is optimal and that one can have high confidence in inferring the presence of the particular body fluid. However if the biomarker is highly specific but an extremely very low expresser then it might be worth considering finding a suitable replacement with more robust expression for future iterations of the current assay.

A simple graphic method was developed to permit categorical inference for the presence of a particular body fluid. The primary raw data from the assay for each sample comprises read counts from each of the 33 biomarkers. Like any bio-analytical system analytical noise is present and the objective of the assay is to distinguish signal from noise. Here we take a pragmatic ad hoc approach to removing sequencing noise by applying a number of filters to the raw read count data at the sample level and the individual biomarker level within each sample. The read count data is then normalized by calculating the relative expression frequencies of each of the biomarkers in the sample expressed as a percentage. Percentage expression frequencies of biomarkers from the same body fluid class are added together such that each sample is displayed as a bar graph showing the relative contributions of each of the six possible biomarker classes (blood, semen, saliva, vaginal secretions, menstrual blood and skin). The noise filter thresholds will need to be fine-tuned as more data from more samples are analyzed but are not expected to be wildly different to what we use at present.

An alternative, complementary body fluid inference method that uses the raw count data unfiltered for noise that we also have employed is agglomerative hierarchical clustering. This method calculates the similarities and differences in biomarker expression between samples and clusters them according to which samples are more similar to one another. Single source samples tested with the targeted RNA assay and subjected to clustering analysis separates the samples and produces clusters that are found to be body fluid specific (Figures 1 and 4). In order to infer the presence of a particular body fluid, several control samples that represent all 6 of the body fluids and skin (4-5 biological replicates per body fluid/tissue) plus any unknown samples are then co-tested with the targeted RNA assay. Agglomerative hierarchical clustering is then carried out with the body fluid controls plus an unknown sample to determine which cluster it belonged. If the unknown sample clusters with a particular body fluid and the correlation of gene expression is high with other markers within the cluster then a categorical inference can be made.

Apart from these ad hoc categorical inference methods, other methods that assign probabilities to predictions are possible. The development and use of a promising probabilistic approach that determines posterior probabilities for each of the possible body fluids will be the subject of a subsequent manuscript.

The results from the various mixture types tested demonstrates the ability of the assay to detect two fluids in binary admixed samples, but also demonstrates the variability that is observed with minor component fluids in some mixtures, especially when the minor component is saliva which has more low- to moderate- expressing biomarkers than the other body fluids. Future work employing more extensive mixture sample sets will be necessary to establish appropriate thresholds for inferring the presence of contributor body fluid types in binary and more complex mixtures.

Despite the ability to definitively identify the body fluids present in a mixture, it is not possible to associate the component DNA profiles with specific body fluids, a requirement in order to meet the goal of obtaining probative objective ‘activity level’ information in investigations. Future developments of the current assay will employ coding region SNPs (or RNA-SNPs) judiciously chosen to be present in the body fluid specific mRNA genes already targeted. This should permit an association of a DNA profile with a specific body fluid or tissue in admixed samples.

## Acknowledgements

This work was supported by EUROFORGEN-NoE, FP7/2007-2013, grant agreement no. 285487 and the National Institute of Justice (NIJ), Office of Justice Programs, U.S. Department of Justice (Award No. 2014-DN-BX-K019). The funding agencies had no role in study design, data analysis and interpretation and in manuscript preparation and submission. The opinions, findings and conclusions or recommendations are those of the authors and do not necessarily reflect those of the funding agencies.

---

**Supplementary Table 1.**
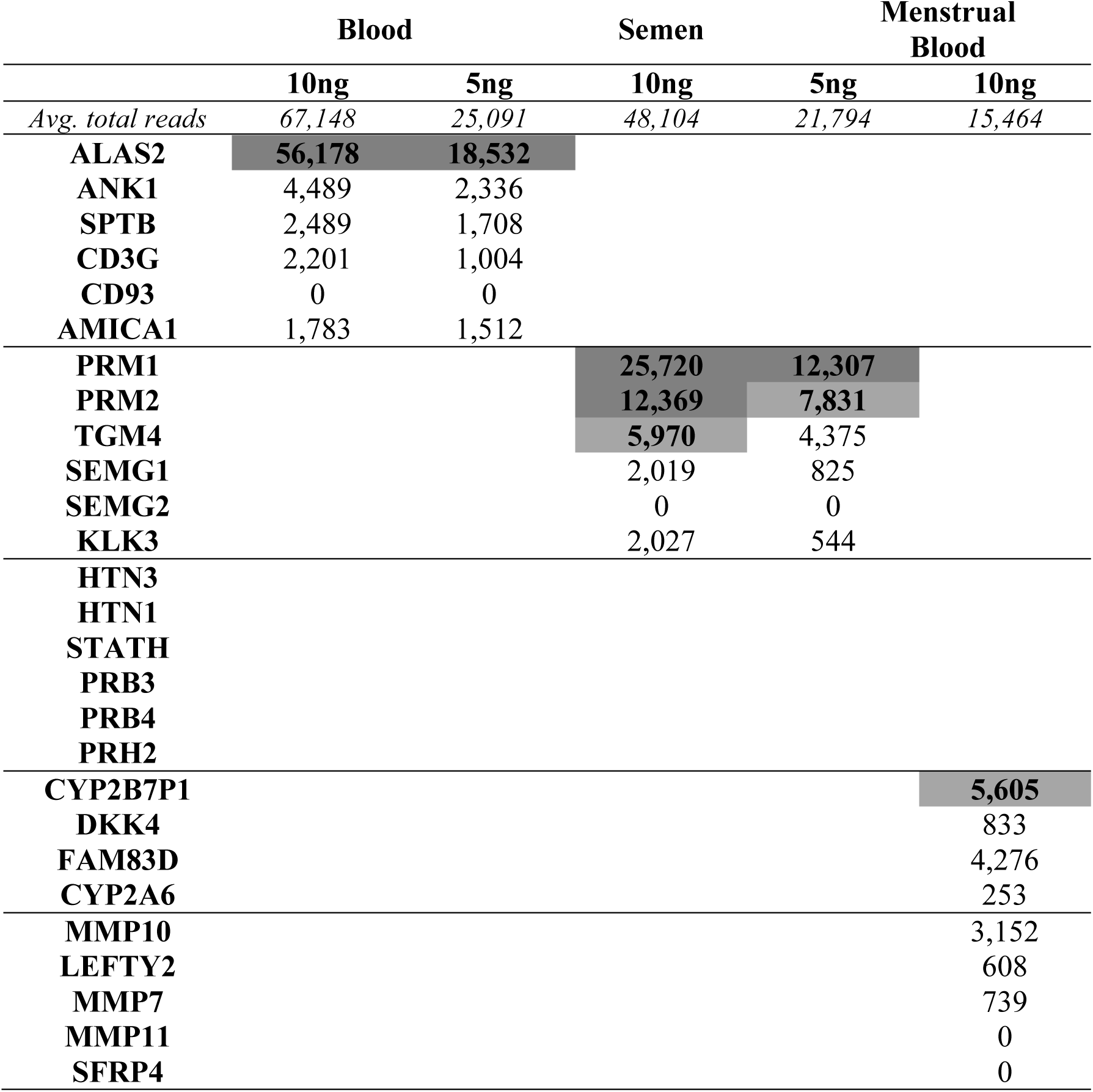
Biomarker Sensitivity. Average total counts per biomarker shown for blood, semen and menstrual blood samples using 5 – 10 ng inputs (N=2 for each input amount). Shading: dark grey ≥ 10,000 read counts; light grey 5,001 – 9,999 read counts; no color ≤5,000 read counts.

**Supplementary Table 2.**
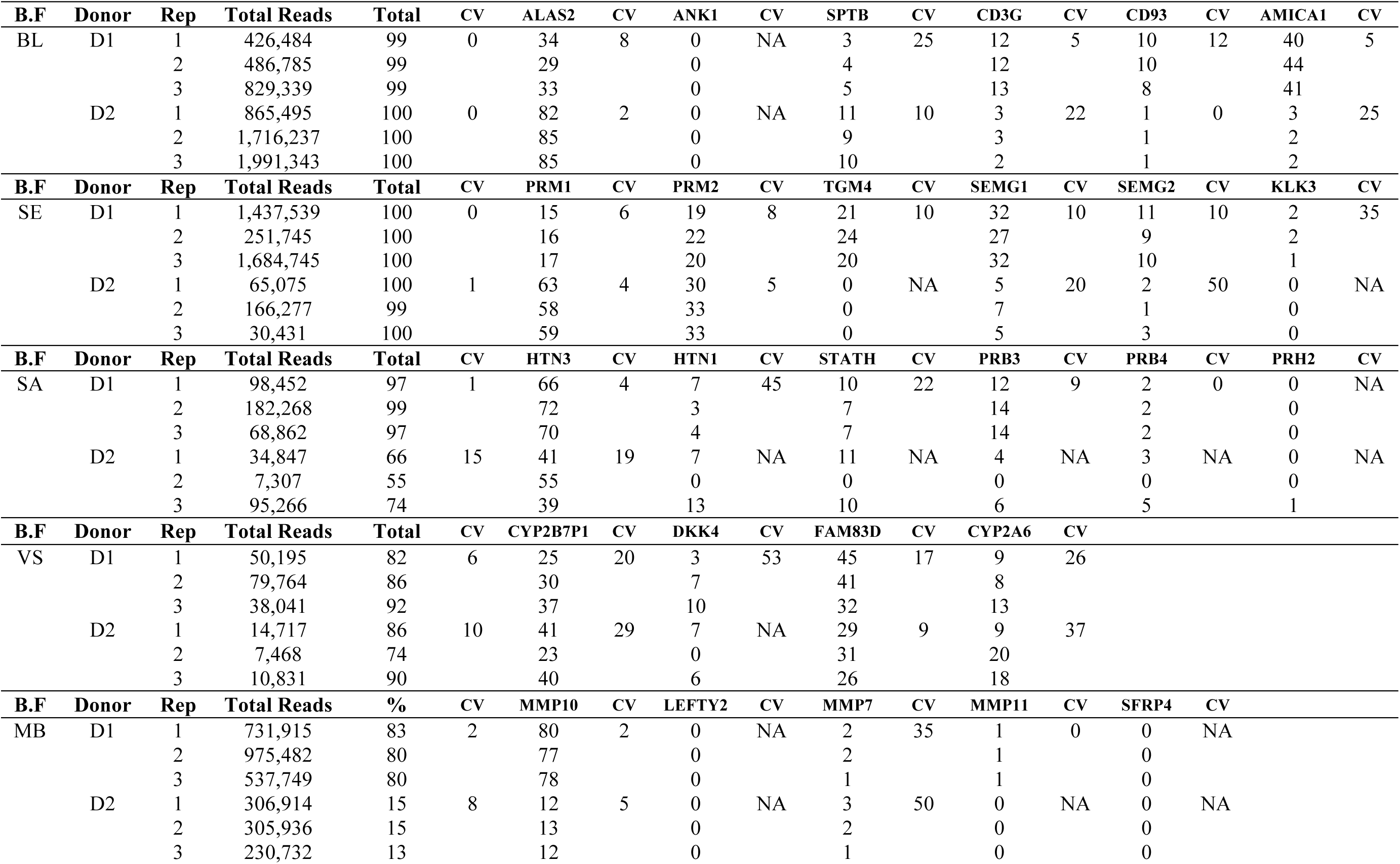
Repeatability of Targeted RNA Sequencing Assay. Percent contributions (biomarker reads/total reads*100) provided under each biomarker. CV (coefficient of variation) was calculated for each triplicate set. NA = CV not determined since triplicate data not available (one or more replicate with value of 0). B.F = Body fluid; Total = percent contribution of target biomarker class for each sample. BL= blood, SE =semen, SA = saliva, VS= vaginal secretions, MB = menstrual blood

**Supplementary Table 3.** Read Counts for Reproducibility Study. B.F = Body fluid; Read counts provided for each biomarker; ng = input ng in assay

| B.F | Donor | ng | Rep | Total Reads | ALAS2 | ANK1 | SPTB | CD3G | CD93 | AMICA1 |
| --- | --- | --- | --- | --- | --- | --- | --- | --- | --- | --- |
| BL | D1 | 41 | 1 | 426,484 | 144,639 | 0 | 13,103 | 52,865 | 40,953 | 172,525 |
|  |  |  | 2 | 486,785 | 139,909 | 0 | 20,909 | 58,914 | 48,626 | 215,017 |
|  |  |  | 3 | 829,339 | 273,790 | 0 | 39,393 | 109,460 | 68,829 | 337,867 |
|  | D2 | 34 | 1 | 865,495 | 703,537 | 0 | 98,017 | 30,000 | 7,774 | 26,167 |
|  |  |  | 2 | 1,716,237 | 1,455,585 | 0 | 156,338 | 51,851 | 11,837 | 40,626 |
|  |  |  | 3 | 1,991,343 | 1,684,303 | 0 | 196,372 | 46,551 | 18,222 | 45,895 |
| B.F | Donor |  | Rep | Total Reads | PRM1 | PRM2 | TGM4 | SEMG1 | SEMG2 | KLK3 |
| SE | D1 | 20 | 1 | 1,437,539 | 213,087 | 278,975 | 308,480 | 454,651 | 153,976 | 28,370 |
|  |  |  | 2 | 251,745 | 39,188 | 54,733 | 61,080 | 67,398 | 23,827 | 5,519 |
|  |  |  | 3 | 1,684,745 | 280,243 | 334,463 | 342,534 | 546,361 | 161,580 | 19,556 |
|  | D2 | 13 | 1 | 65,075 | 41,360 | 19,449 | 0 | 3,280 | 986 | 0 |
|  |  |  | 2 | 166,277 | 96,489 | 55,864 | 0 | 11,116 | 1,037 | 0 |
|  |  |  | 3 | 30,431 | 17,816 | 10,416 | 0 | 1,420 | 779 | 0 |
| B.F | Donor |  | Rep | Total Reads | HTN3 | HTN1 | STATH | PRB3 | PRB4 | PRH2 |
| SA | D1 | 20 | 1 | 98,452 | 64,901 | 7,223 | 9,256 | 11,610 | 2,031 | 0 |
|  |  |  | 2 | 182,268 | 131,648 | 5,720 | 13,598 | 26,053 | 4,167 | 0 |
|  |  |  | 3 | 68,862 | 48,000 | 3,014 | 4,999 | 9,662 | 1,170 | 0 |
|  | D2 | 76 | 1 | 34,847 | 14,368 | 2,434 | 3,830 | 1,517 | 946 | 0 |
|  |  |  | 2 | 7,307 | 4,047 | 0 | 0 | 0 | 0 | 0 |
|  |  |  | 3 | 95,266 | 37,489 | 12,754 | 9,201 | 5,689 | 4,893 | 835 |
| B.F | Donor |  | Rep | Total Reads | CYP2B7P1 | DKK4 | FAM83D | CYP2A6 |  |  |
| VS | D1 | 79 | 1 | 50,195 | 12,396 | 1,275 | 22,653 | 4,476 |  |  |
|  |  |  | 2 | 79,764 | 23,998 | 5,317 | 32,392 | 6,612 |  |  |
|  |  |  | 3 | 38,041 | 14,110 | 3,750 | 11,989 | 5,052 |  |  |
|  | D2 | 66 | 1 | 14,717 | 6,017 | 980 | 4,306 | 1,369 |  |  |
|  |  |  | 2 | 7,468 | 1,713 | 0 | 2,292 | 1,526 |  |  |
|  |  |  | 3 | 10,831 | 4,370 | 620 | 2,871 | 1,926 |  |  |
| B.F | Donor |  | Rep | Total Reads | MMP10 | LEFTY2 | MMP7 | MMP11 | SFRP4 |  |
| MB | D1 | 63 | 1 | 731,915 | 588,308 | 0 | 16,392 | 3,914 | 0 |  |
|  |  |  | 2 | 975,482 | 749,713 | 0 | 19,131 | 10,585 | 0 |  |
|  |  |  | 3 | 537,749 | 415,722 | 0 | 7,368 | 5,046 | 0 |  |
|  | D2 | 57 | 1 | 306,914 | 36,217 | 0 | 9,586 | 0 | 0 |  |
|  |  |  | 2 | 305,936 | 39,852 | 0 | 7,290 | 0 | 0 |  |
|  |  |  | 3 | 230,732 | 27,682 | 0 | 1,725 | 0 | 0 |  |

**Supplementary Table 4.**
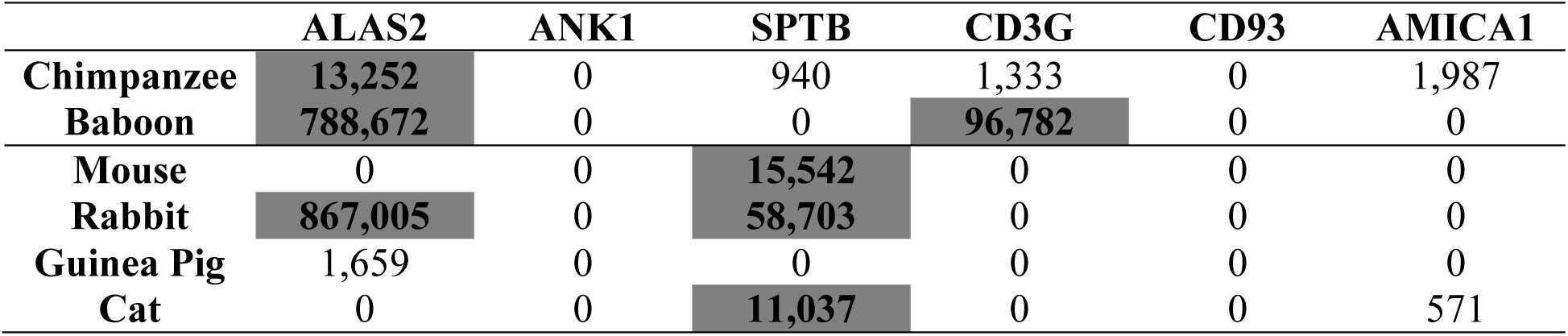
Species Specificity – Blood Biomarkers. Read counts of each biomarker (N=1 for each species). Shading: dark grey ≥ 10,000 read counts; no color ≤5,000 read counts

**Supplementary Table 5.** Specificity - Tissues. Read counts of each biomarker (N=1 for each tissue). Shading: dark grey ≥ 10,000 read counts; no color ≤5,000 read counts.

| Body Fluid | Biomarker | Brain | Small Intestine | Trachea | Liver | Skeletal Muscle |
| --- | --- | --- | --- | --- | --- | --- |
| BL | ALAS2 |  |  |  | 1,465 |  |
|  | ANK1 | 21,521 |  |  |  | 4,872 |
|  | SPTB | 13,263 |  |  | 1,129 | 11,112 |
|  | CD3G |  | 1,781 |  |  |  |
|  | CD93 | 981 | 1,419 |  |  |  |
|  | AMICA1 | 1,607 | 897 |  | 1,645 |  |
| SE | PRM1 |  |  |  | 3,594 | 1,110 |
|  | PRM2 |  |  |  | 4,527 | 542 |
| SA | STATH |  |  | 1,514 |  |  |
|  | PRB3 |  |  | 45,465 |  |  |
|  | PRB4 |  |  | 11,266 |  |  |
|  | PRH2 |  |  | 3,698 |  |  |
| VS | CYP2B7P1 |  | 508 |  |  |  |
|  | FAM83D | 1,847 |  |  |  |  |
| MB | MMP7 | 629 |  |  |  |  |
| SK | CCL27 | 674 |  |  |  |  |

*\*Lung, heart, kidney, adipose and stomach were tested but did not meet 5,000 minimum total read count threshold*

**Supplementary Table 6.**
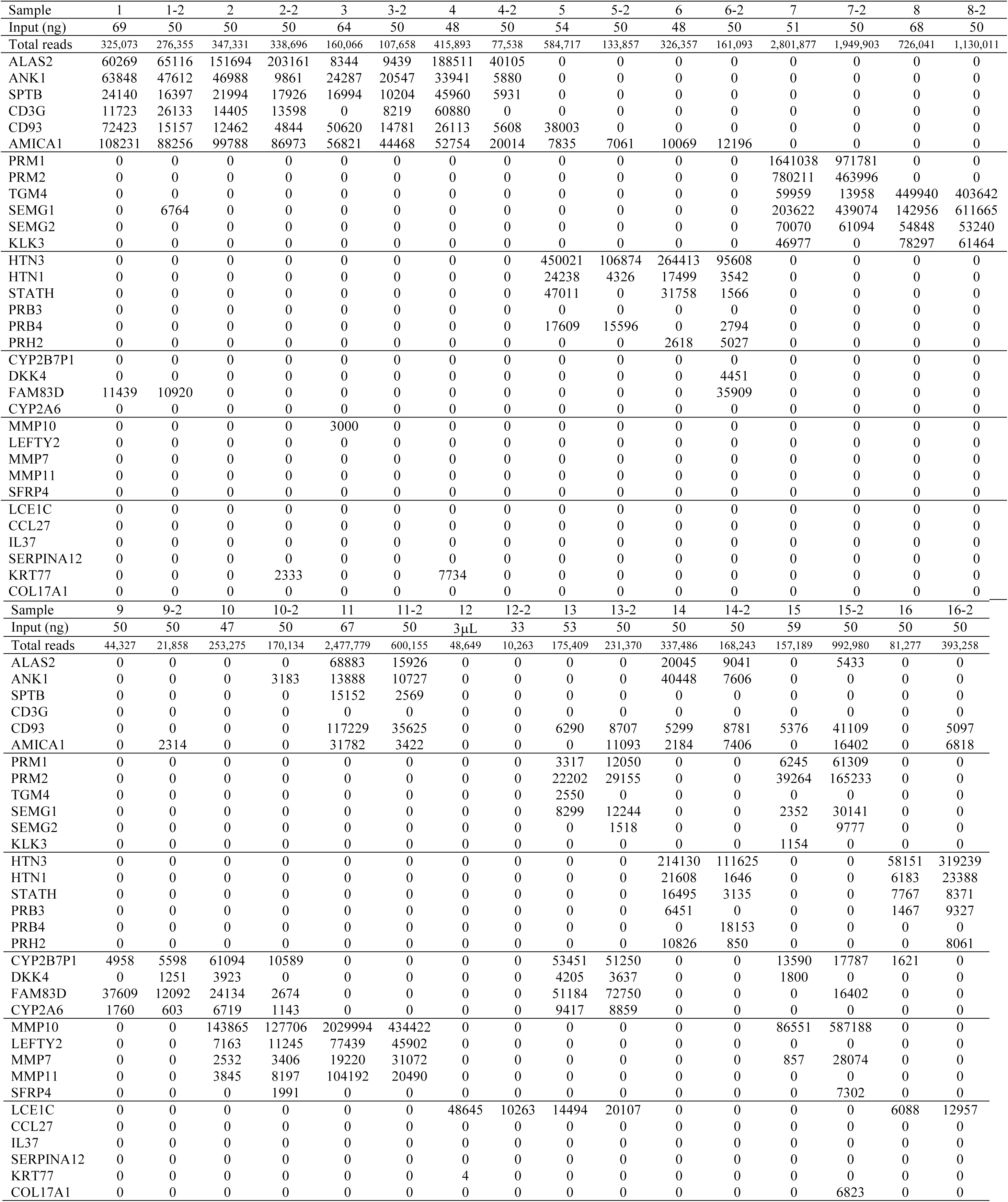
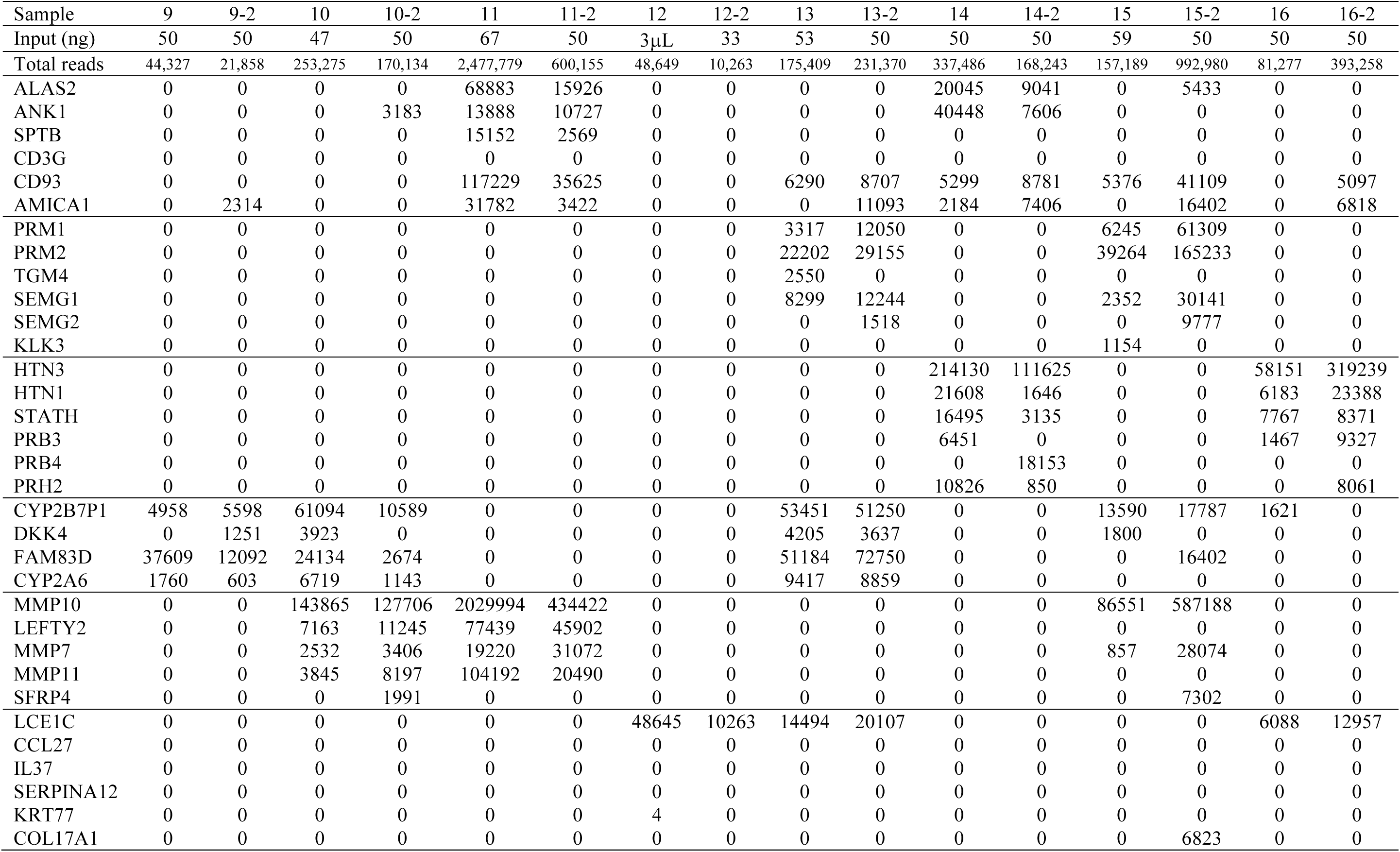
Read Counts for 32 Samples In Blind Study. -2 samples = duplicate set tested by organizers of the blind study (Lab 2)

## Supplementary Figure Legends

**Supplementary Figure 1.**
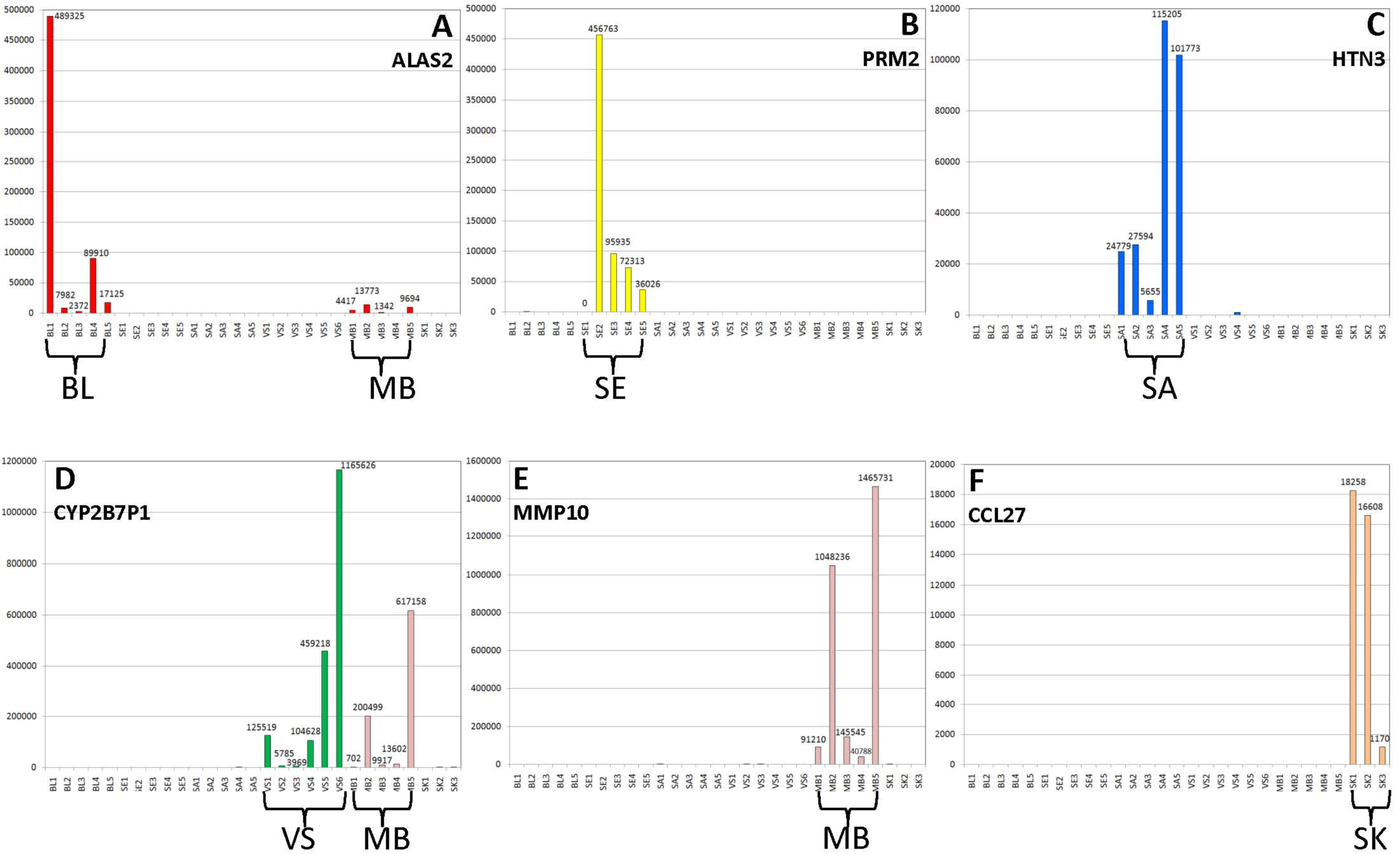
Body Fluid Specific Gene Expression Exemplified by Individual Gene Candidates amongst 29 Body Fluid Samples. Read counts for individual biomarkers (A – ALAS2, blood-specific; B – PRM2, semen (i.e. sperm)-specific; C – HTN3, saliva-specific; D – CYP2B7P1, vaginal-specific; E – MMP10, menstrual blood-specific; F – CCL27, skin-specific) are shown amongst a set of 29 body fluid samples (BL=blood (N=5), SE=semen (N=5), SA=saliva (N=5), VS=vaginal (N=6), MB=menstrual blood (N=5), SK=skin (N=3). Colored bars represent biomarker expression (i.e. read counts) in the target body fluid: blood (red), semen (yellow), saliva (blue), vaginal (green), menstrual blood (pink) and skin (peach). For full reference to colors, readers are directed to the online version of the article. Y-axis – read counts, X-axis – body fluid samples.

**Supplementary Figure 2.**
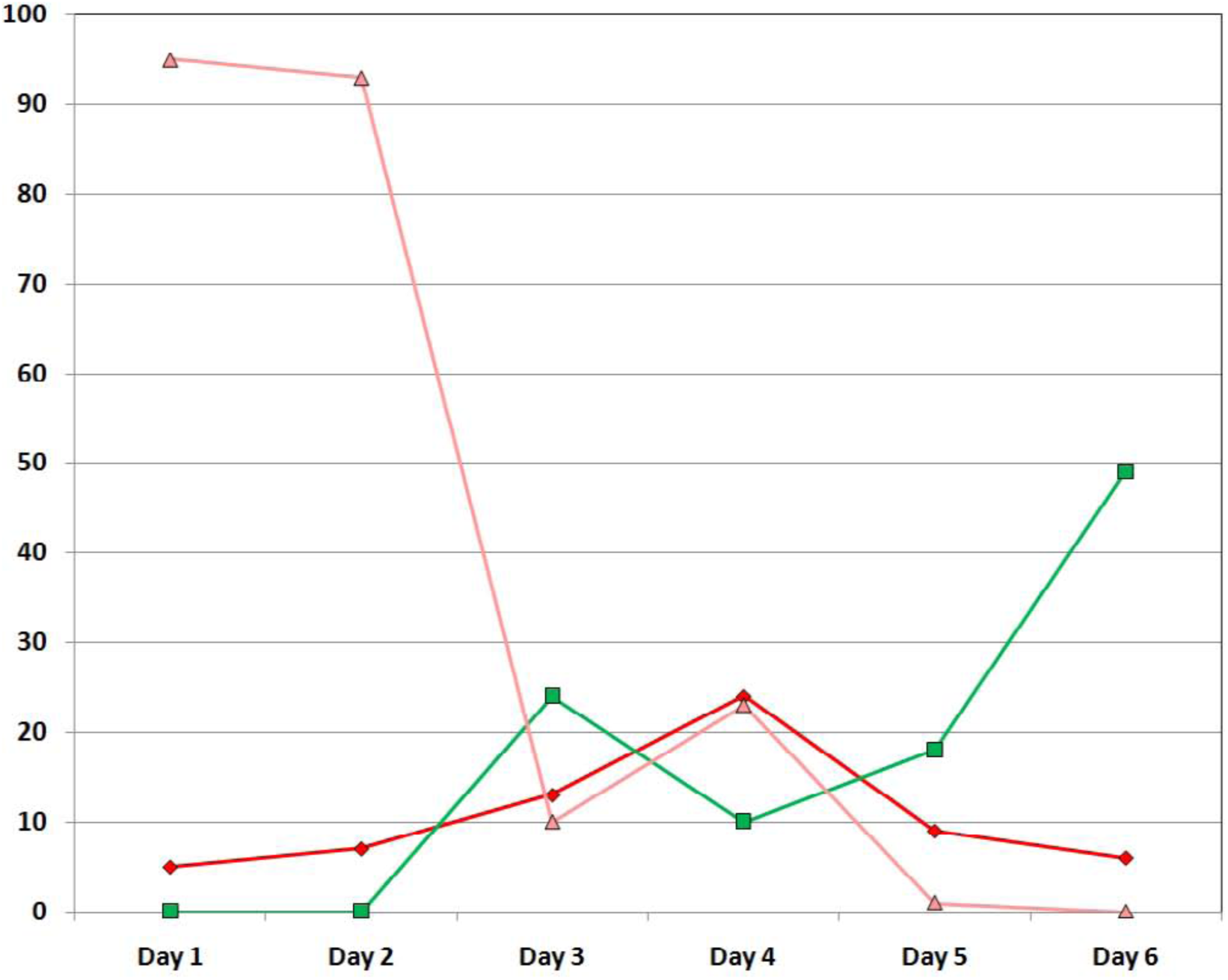
Menstrual Blood Biomarker Class Expression over 6-day Menstruation Cycle. The percent contribution for individual biomarkers was calculated (reads per biomarker/total reads per sample). The percentage reads from each body fluid specific biomarker were combined into their respective biomarker classes in order to determine the percentage of reads per sample attributable to each body fluid or tissue class. Percent reads attributable to each biomarker class are diagrammed and represented by color: red – expression from blood biomarkers, green – expression from vaginal secretions biomarkers and pink – expression from menstrual blood biomarkers. Percent composition is plotted for each biomarker class for each sample collected every day during a 6-day period of reported menstruation. Y-axis–percent contribution; X-axis – day of menstruation.

**Supplementary Figure 3.**
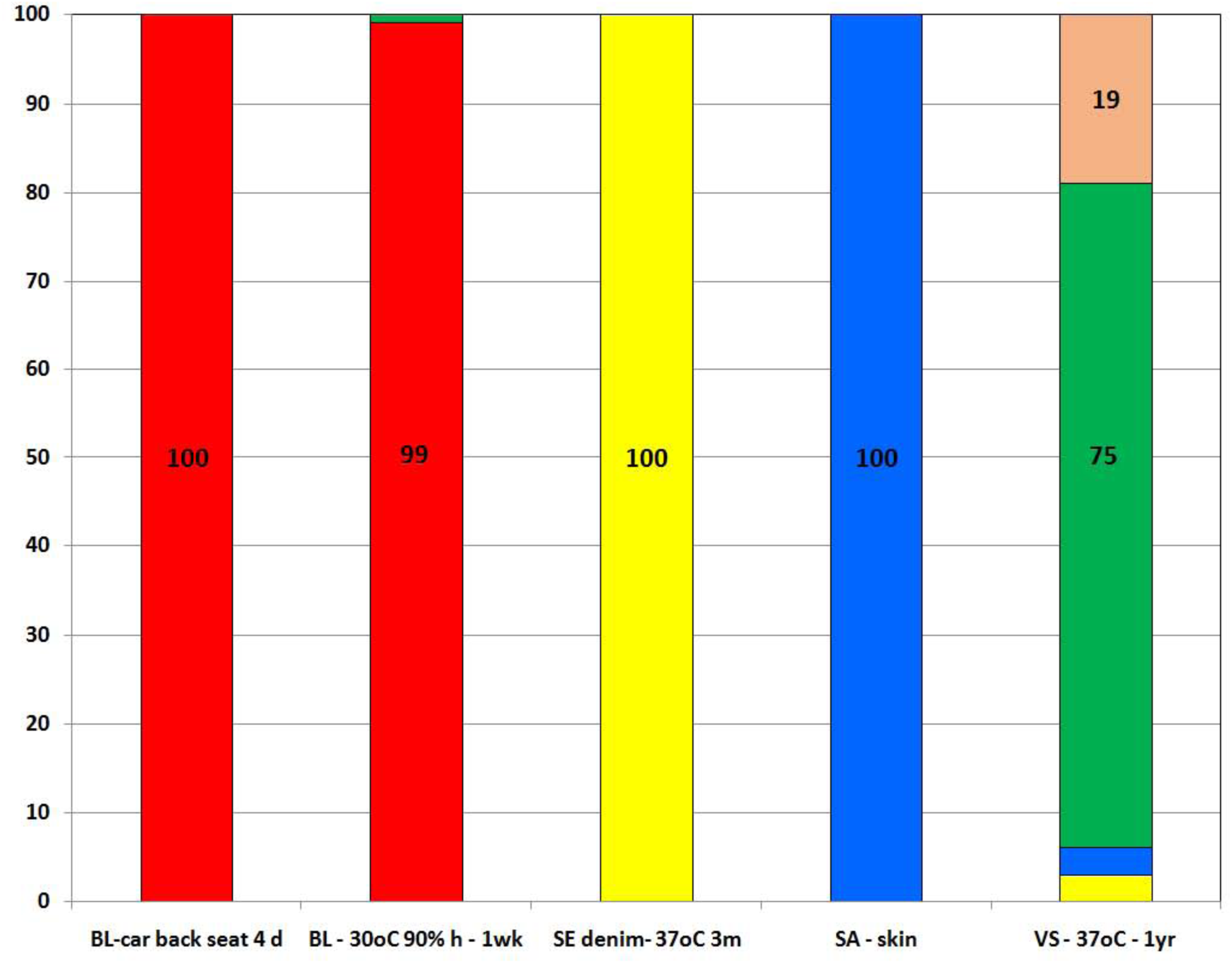
Biomarker Class Expression Composition in Mock Casework and Environmentally Compromised Body Fluid Samples. Body fluid samples were exposed to various storage temperatures and humidity levels: blood in the back seat of a car for 4 days (BL- car back seat 4d), blood stored at 30°C and 90% humidity for 1 week (BL-30°C 90% h – 1 wk), semen on denim incubated at 37°C for 3 months (SE denim 37°C 3 m) and vaginal secretions incubated at 37°C for 1 year (VS – 37°C – 1 yr). Additionally saliva was placed directly on skin surface (arm) (SA – skin). For each sample, the percentage of reads per sample attributable to each body fluid or tissue class was determined. Percent reads attributable to each biomarker class are listed and represented by color: red – expression from blood biomarkers, yellow – expression from semen biomarkers, blue – expression from saliva biomarkers, green – expression from vaginal secretions biomarkers and peach – expression from skin biomarkers. Y-axis – percent contribution; X-axis – sample.

